# Astrocyte-Secreted Neurocan Controls Inhibitory Synapse Formation and Function

**DOI:** 10.1101/2023.04.03.535448

**Authors:** Dolores Irala, Shiyi Wang, Kristina Sakers, Leykashree Nagendren, Francesco Paolo Ulloa-Severino, Dhanesh Sivadasan Bindu, Cagla Eroglu

## Abstract

Astrocytes strongly promote the formation and maturation of synapses by secreted proteins. To date, several astrocyte-secreted synaptogenic proteins controlling different stages of excitatory synapse development have been identified. However, the identities of astrocytic signals that induce inhibitory synapse formation remain elusive. Here, through a combination of *in vitro* and *in vivo* experiments, we identified Neurocan as an astrocyte-secreted inhibitory synaptogenic protein. Neurocan is a chondroitin sulfate proteoglycan that is best known as a protein localized to the perineuronal nets. However, Neurocan is cleaved into two after secretion from astrocytes. We found that the resulting N- and C-terminal fragments have distinct localizations in the extracellular matrix. While the N-terminal fragment remains associated with perineuronal nets, the Neurocan C-terminal fragment localizes to synapses and specifically controls cortical inhibitory synapse formation and function. Neurocan knockout mice lacking the whole protein or only its C-terminal synaptogenic region have reduced inhibitory synapse numbers and function. Through super-resolution microscopy and *in vivo* proximity labeling by secreted TurboID, we discovered that the synaptogenic domain of Neurocan localizes to somatostatin-positive inhibitory synapses and strongly regulates their formation. Together, our results unveil a mechanism through which astrocytes control circuit-specific inhibitory synapse development in the mammalian brain.

## INTRODUCTION

The development of a properly functioning brain depends on the balanced establishment, coordinated maturation, and activity of excitatory and inhibitory synapses^1^. Therefore, understanding cellular and molecular mechanisms underlying the formation of excitatory and inhibitory synapses and how these processes are orchestrated during development is crucial.

In the mammalian cerebral cortex, 80% of neurons are glutamatergic (excitatory) pyramidal neurons, and 20% are GABAergic (inhibitory) interneurons^2,3^. There is a wide diversity of interneurons based on their morphological, transcriptomic, and electrophysiological properties ^4–6^. The three major cortical interneuron types express Somatostatin (SST), Parvalbumin (PV), or serotonin receptor 3A (Htr3a), and their distribution varies across different cortical areas^2^.

Astrocytes, the major perisynaptic macroglial cells in the brain, actively control synapse formation and function ^7–9^. Previous studies revealed that astrocytes strongly promote excitatory synapse development through the secretion of a variety of synaptogenic proteins, such as thrombospondins, Sparcl1/hevin, glypicans, and chordin-like1^10–13^. It has long been known that astrocytes also control inhibitory synapse formation and maturation via secreted proteins^14,15^. Yet, the identities of these astrocyte-secreted inhibitory synaptogenic signals and whether astrocytes control the formation of all types of inhibitory synapses are still unknown.

Astrocyte-secreted excitatory synaptogenic factors were previously identified using a purified retinal ganglion cell (RGC) neuron-only culture system ^16^. In this system, postnatal neurons from rat or mouse retinas are isolated via immunopanning, resulting in a nearly pure population of RGCs. These neurons survive under serum-free conditions supported by defined growth factors; however, they form very few synapses amongst each other when cultured alone. The addition of astrocyte-conditioned media (ACM) robustly promotes the formation of synaptic connections between neurons^17,18^. Therefore, this neuronal system allowed for the identification of synaptogenic ACM proteins, such as thrombospondins^10^, Sparcl1/hevin^11^, Glypicans^12^, Chdl1^13^. Moreover, using this system, the neuronal mechanisms underlying astrocyte-induced excitatory synaptogenesis were deciphered^19–21^. However, because RGCs are glutamatergic neurons, they could not be used to study the role of astrocytes in inhibitory synapse formation.

Previously, embryonic hippocampi were used to generate glia-depleted cultures containing inhibitory and excitatory neurons^14,15^. Treatment of these embryonic neuron cultures with ACM induced inhibitory synaptogenesis, indicating that astrocytes control both excitatory and inhibitory synapse formation^14,15^. Interestingly, the astrocytic factors that induce excitatory synaptogenesis do not induce inhibitory synapse formation^15^, suggesting that astrocytes must promote inhibitory synaptogenesis through distinct pathways. However, in these hippocampal cultures, because neurons were isolated from embryos, it has been hard to distinguish the effects of astrocytic factors on neuronal maturation from their roles in synapse formation. Thus, the identities of the astrocytic factors controlling inhibitory synapse formation remained unknown for more than a decade.

To eliminate this technical bottleneck, here we developed an *in vitro* system using immuno-panned postnatal cortical neurons. This system is suitable for studying the roles of astrocytes in inhibitory synapse formation because, like the rodent cortex, it is composed of ~80% excitatory neurons and 20% inhibitory neurons, and neurons can be cultured under serum-free conditions in the absence of glia. We found that treating these cortical neurons with ACM robustly induces the formation of excitatory and inhibitory synapses without affecting neuronal survival, providing an optimal system to identify inhibitory synaptogenic proteins secreted by astrocytes.

Using this system, we identified the chondroitin sulfate proteoglycan Neurocan (NCAN) as an astrocyte-secreted molecule that robustly induces inhibitory synaptogenesis in the cerebral cortex. NCAN is part of the Lectican family of proteins (Aggrecan, Versican, Neurocan, and Brevican). Lecticans are well known as one of the main components of Perineuronal Nets, an extracellular matrix structure that surrounds neuronal cell bodies and controls synaptic plasticity^22,23^. In particular, *NCAN* mutations are a risk factor for bipolar disorder (BD), Schizophrenia (SCZ), and mania^24–27^. NCAN is composed of three main parts: an N-terminal composed of an Ig-module and two Link domains, which are known to be located in PNNs through hyaluronic acid, a central linker region that carries the glycosaminoglycan (GAG) chains, and a C-terminal region composed of two EGF-like domains, a lectin-like domain, and a sushi domain which might be involved in controlling cell adhesion^28^. Interestingly, NCAN is cleaved during postnatal development into two fragments of roughly the same size, the NCAN N-terminal and C-terminal ^28^. The role of the NCAN N-terminal during brain development has been widely studied. It is well known that it interacts with hyaluronic acid through its link domains, and it is involved in PNN formation and function ^29,30^. Unbound NCAN N-terminal acts as an inhibitor of cell-adhesion molecules. For example, it inhibits Semaphorin 3F-induced spine remodeling, NCAM/EphA3 mediated axonal repulsion, and β1-Integrin–mediated adhesion and neurite outgrowth ^31–33^. NCAN C-terminal is also abundant in the ECM, and it contains multiple protein domains associated with cell adhesion; however, the role of this fragment in the CNS remains underexplored.

Here, by generating two new NCAN mutant mice and using a combination of super-resolution, electron microscopy, and electrophysiology approaches, we found that NCAN controls the formation and function of inhibitory synapses in the developing mouse cortex through interactions mediated by its C-terminal ELS-domains. Using proteomic labeling by TurboID-tagged NCAN domains, we identified that the synaptogenic ELS domain interacts with multiple inhibitory synaptic proteins, including the neuropeptide somatostatin (SST), which is specifically expressed by the SST+ interneurons and presynaptically released. Importantly, in glia-free neuronal cultures, NCAN ELS domain treatment specifically induced SST+ inhibitory synaptogenesis, and SST+ synapse numbers and function were severely reduced in transgenic mice lacking the NCAN ELS domain. Our results shed light on the role of astrocytes during inhibitory synapse formation and provide a circuit-specific mechanism through which astrocytes control inhibition.

## RESULTS

### NCAN induces inhibitory synaptogenesis in glia-free cortical neuron cultures

Astrocytes potently induce synapse formation through secreted proteins ^10–12^. In the past two decades, several synaptogenic proteins that induce the formation and functional maturation of excitatory synapses have been identified; however, the identity of the astrocyte-secreted proteins controlling inhibitory synaptogenesis remained elusive. To address this knowledge gap, we optimized a rodent glia-free cortical neuron culture system which includes both excitatory and inhibitory neurons. To do so, neurons were isolated from neonatal rat cortices by L1CAM immunopanning and were cultured in the absence of serum but in the presence of well-described growth factors (see methods for details). These neurons survived over two weeks in culture and developed extensive neurites even in the absence of glial-feeder layers. We also isolated astrocytes from neonatal rat cortices to prepare astrocyte-conditioned media (ACM), as described before ^34^. To determine the synaptogenic effects of ACM or recombinant proteins on the purified neurons, we treated them with ACM or recombinant proteins between at days *in vitro* (DIV) 8 and 11 (Fig 1A).

**Figure 1:**
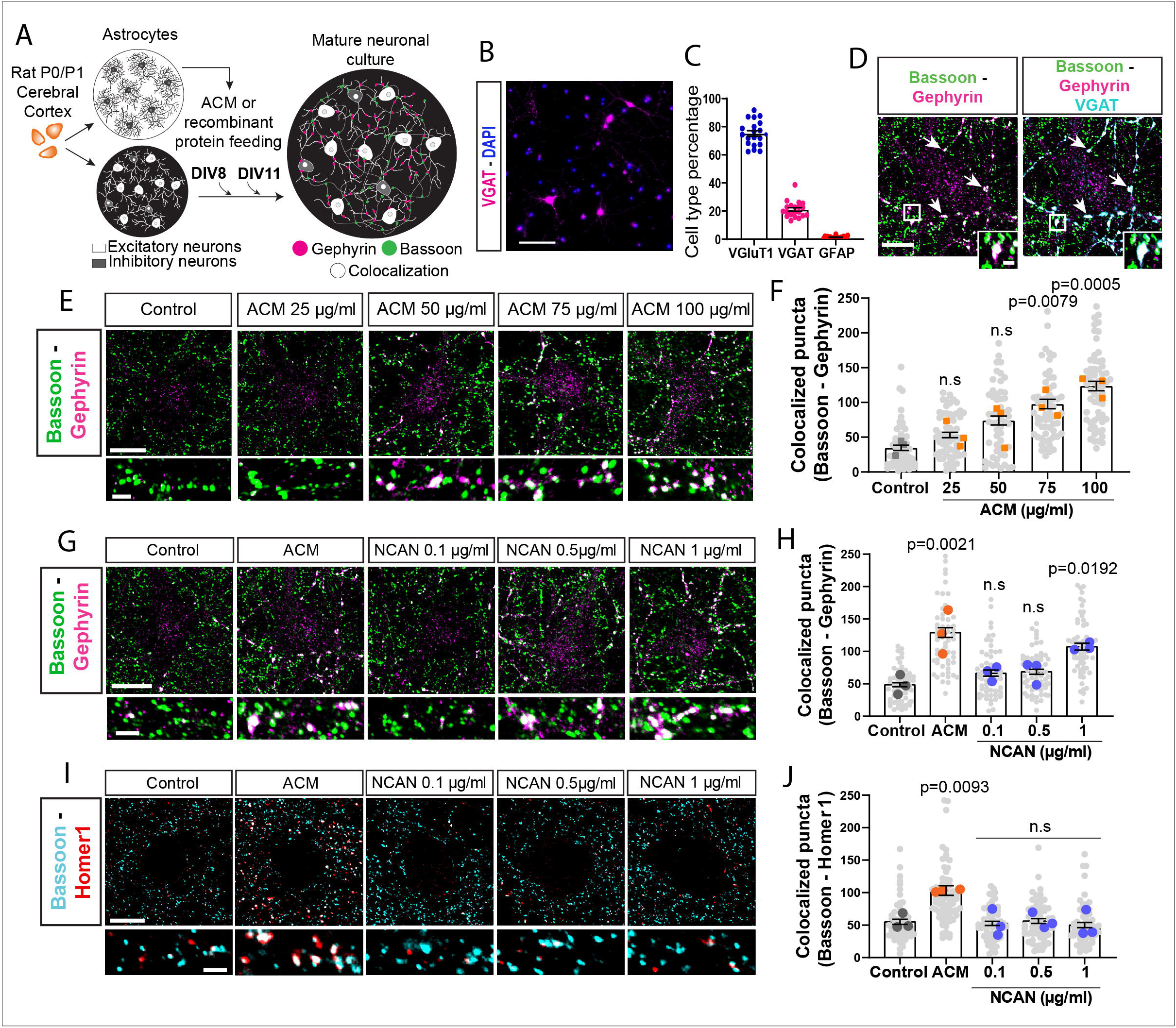
NCAN induces inhibitory synapse formation in postnatal glia-free neuronal cultures. (A) Schematic of neuronal culture assay and feeding schedule. (B) Representative image of inhibitory GABAergic interneurons labeled with GABAergic neuron marker VGAT (magenta) in glia-free neuronal cultures. Scale bar, 50μm. C) Percentage of cell types in glia-free neuronal cultures. Excitatory glutamatergic neurons (VGluT1+), inhibitory GABAergic neurons (VGAT+), and astroglia (GFAP+). Data are mean ± s.e.m. 20 images/condition. (D) Representative images of inhibitory synapses marked with presynaptic marker Bassoon (green) and VGAT (cyan) together with postsynaptic marker Gephyrin (magenta) in glia-free neuronal cultures. Scale bar, 15μm. Insert scale bar: 2μm. (E) Dose-response curve of astrocyte-conditioned media in glia-free neuronal cultures. Inhibitory synapses were marked as the apposition of the presynaptic marker Bassoon and postsynaptic marker Gephyrin. Scale bar: 20μm. Insert scale bar: 5μm. (F) Quantification of inhibitory synapses with increasing concentrations of ACM. Synapses are identified as co-localization of Bassoon and Gephyrin signals. Data are mean ± s.e.m. n = 3 independent experiments, 20 cells/condition/experiment. One-way ANOVA, Dunnett’s post-test. (G) Dose-response curve of increasing concentration of NCAN recombinant protein in glia-free neuronal cultures. Inhibitory synapses were marked with presynaptic marker Bassoon and postsynaptic marker Gephyrin. Scale bar: 25μm. Insert scale bar: 5μm. (H) Quantification of inhibitory synapses with increasing concentrations of NCAN recombinant protein. Data are mean ± s.e.m. n = 3 independent experiments, 20 cells/condition/experiment. One-way ANOVA, Dunnett’s post-test. (I) Dose-response curve of increasing concentration of NCAN recombinant protein in glia-free neuronal cultures. Excitatory synapses were marked with presynaptic marker Bassoon (cyan) and postsynaptic marker Homer1 (red). Scale bar: 25μm. Insert scale bar: 5μm. (J) Quantification of excitatory synapses with increasing concentrations of NCAN recombinant protein. Excitatory synapses were marked as the co-localization of Bassoon and Homer1. Data are mean ± s.e.m. n = 3 independent experiments, 20 cells/condition/experiment. One-way ANOVA, Dunnett’s post-test.

The resulting cortical neuron cultures contained less than 2% GFAP-positive astroglia, 78% excitatory neurons marked with Vesicular Glutamate transporter 1 (VGluT1+), and 20% of inhibitory neurons identified with Vesicular GABA Transporter (VGAT+) staining (Figures 1B and 1C). We identified both SST+ (~9%) and PV+ (~11%) interneurons in these glia-free neuronal cultures at DIV 14 (Figures S1A and S1B).

Using this culture system, we studied how inhibitory neurons (identified as VGAT^+^) make synaptic contacts with excitatory neurons (identified as VGAT^-^). To visualize inhibitory synapses, we labeled the synaptic contacts with presynaptic markers Bassoon (active zone) and VGAT (inhibitory synaptic vesicles and neurites) together with the postsynaptic marker Gephyrin (Figure 1D). To determine whether glia-free neuronal cultures increase the number of inhibitory synapses in the presence of astrocyte-secreted factors, we exposed the neurons to rising concentrations of ACM. We found that treating neurons with ACM robustly induces the formation of inhibitory synapses marked as the close apposition of presynaptic Bassoon and postsynaptic Gephyrin onto cortical pyramidal neurons, without altering the numbers of individual Bassoon and Gephyrin puncta *in vitro*. (Figures 1E and 1F). The addition of ACM does not alter the percentage of VGAT+ inhibitory neurons in these cultures (Figures S1C and S1D), indicating that this increase in inhibitory synapse numbers is not due to the enhanced survival of GABAergic neurons.

To identify the astrocyte-secreted proteins that promote inhibitory synapse formation, we conducted a candidate-based screen. To choose candidates, we cross-referenced three proteomic studies that identified proteins present in the ACM^12,35,36^ with a cell-type-specific RNA expression database from the mouse cortex^37^. We selected four candidates: Clusterin (Clu), Alpha-Dystroglycan1 (DAG), Neurocan (NCAN), and Carboxypeptidase E (CpE), because all three proteomic studies identified these candidates to be present in the ACM *in vitro,* and their expression is highly enriched in astrocytes, compared to other CNS cell types *in vivo* (Figure S1E). We tested each candidate’s inhibitory synaptogenic effect *in vitro* using increasing concentrations of each factor. We also tested the effect of TGFβ1 on glia-free neuronal cultures because TGFβ1 was previously proposed as an astrocyte-secreted protein that triggers inhibitory synapse formation in glia-depleted embryonic cortical neuron cultures ^38^. However, the proteomic studies have not detected TGFβ1 in the ACM, and TGFβ1 is predominantly expressed by microglia *in vivo* and not astrocytes ^37^. We found that when added to glia-free postnatal cortical cultures, TGFβ1 did not induce inhibitory synapse formation (Figure S1E).

For our screen, we used 100μg/ml of ACM as our positive control and neuronal growth media as our negative control. We found that DAG, CpE, or Clu do not induce inhibitory synaptogenesis when added to glia-free cortical neuron cultures in any of the tested concentrations (Figures S1F-I). However, 1 μg/ml NCAN robustly induced inhibitory synapse formation, determined by the significant increase in the numbers of co-localized Bassoon and Gephyrin puncta (Figures 1G-H). NCAN did not alter the abundance of Bassoon or Gephyrin puncta (Figures S1J-K). These results suggest that NCAN is an ACM protein that induces inhibitory synapse formation *in vitro*.

Next, we tested whether NCAN can also induce excitatory synaptogenesis *in vitro*. For this analysis, we treated neurons with increasing doses of NCAN and stained them with presynaptic markers Bassoon and Vesicular glutamate transporter 1 (VGluT1, specific to excitatory presynapses) together with postsynaptic marker Homer1 (specific to excitatory synapses). We found that NCAN does not promote an increase in the numbers of excitatory synapses (Bassoon and Homer1 colocalization) made onto excitatory neurons (VGluT1+) *in vitro* at any of the doses used (Figures 1I and 1J). These data reveal that NCAN specifically controls inhibitory but not excitatory synapse formation *in vitro*.

### Astrocytes highly express NCAN that is cleaved into N- and C-terminal fragments during postnatal development

*Ncan* encodes for a large multidomain ECM protein, which is highly glycosylated with long GAG side chains in the middle of the protein (Figure 2A). NCAN is known to be cleaved into two halves generating an N- and C-terminal fragment (Figure 2A)^28^. NCAN fragments contain different protein domains; the N-terminal fragment includes an Ig-module, two Link domains involved in binding hyaluronic acid, and a sequence with multiple GAG-side chains, whereas the C-terminal has GAG-side chains, followed by two EGF-like domains, a lectin-like domain, and a sushi domain (Figure 2A) ^29^. To determine the expression and cleavage of NCAN in the developing mouse brain, we used two antibodies, one recognizing the N-terminus and the other the C-terminus. The N-terminal NCAN antibody specifically recognizes the Ig-module and Link domains (Nter-ab), and the NCAN C-terminal antibody binds to the lectin-like domain (Cter-ab) (see materials and methods) (Figure 2A).

**Figure 2:**
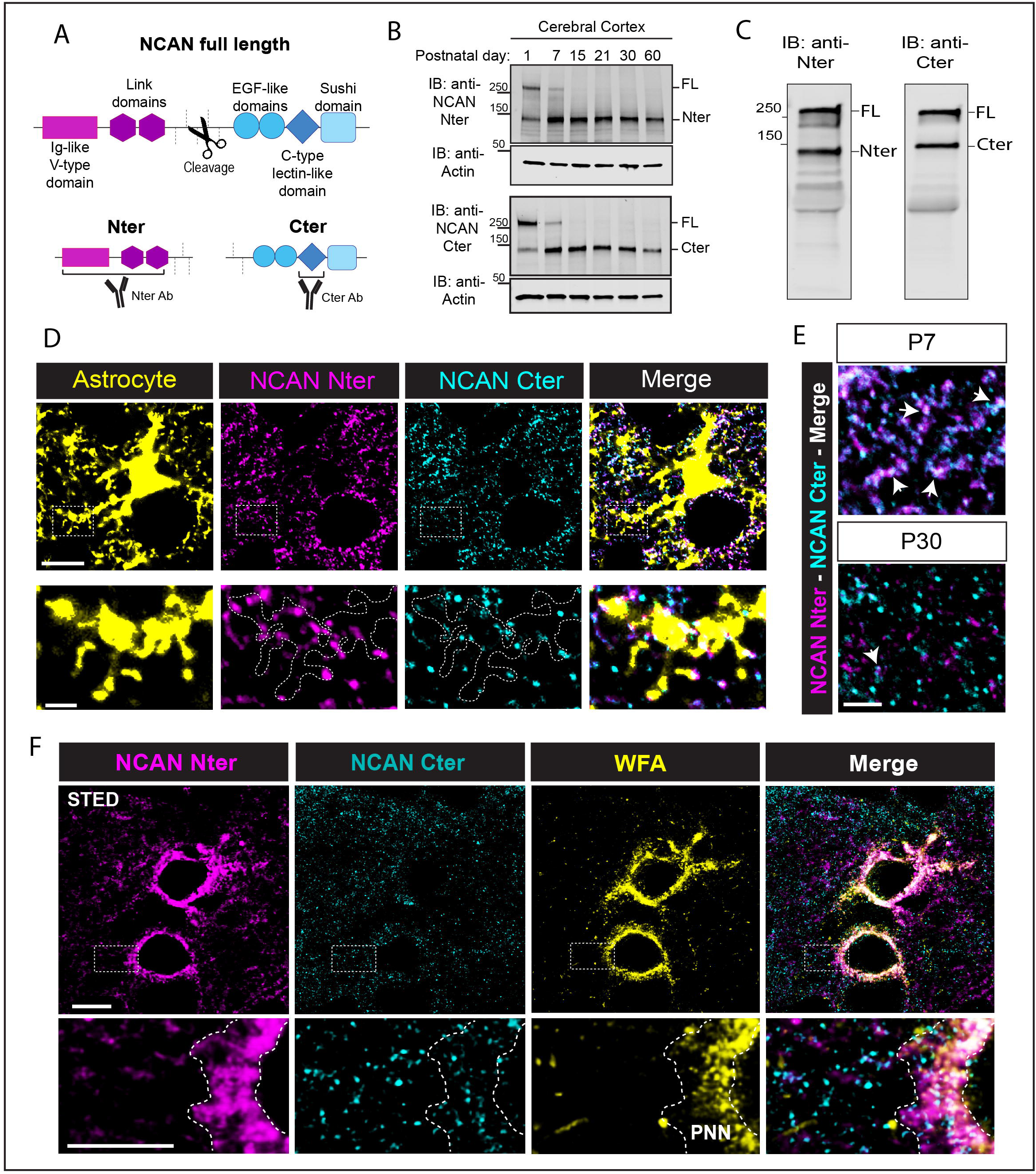
NCAN is expressed by astrocytes during postnatal development and is processed into N and C terminal fragments. (A) Schematic of NCAN full-length protein, cleavage site and resulting NCAN N-terminal (Nter) and C-terminal (Cter) fragments. The antibody epitopes are marked for each fragment. (B) Western blot analysis of NCAN N- and C-terminal expression in postnatal brain cortical lysates at different developmental time points. (C) Western blot analysis of NCAN N- and C-terminal expression in astrocyte-conditioned media. (D) Representative image of NCAN N- (magenta) and C-terminal (cyan) in the cerebral cortex of Aldh1L1-GFP mice at P7. Astrocytes are labeled in yellow. Scale bar: 25μm. Insert scale bar: 5μm. (E) Representative images of NCAN N- and C-terminal distribution in the ECM of WT mice at P7 and P30. Scale Bar: 15μm. (F) Three-color STED image of NCAN N-terminal and C-terminal expression at perineuronal nets (marked in yellow) with WFA. Scale bar: 15μm. Insert scale bar: 10μm

To analyze NCAN expression during postnatal brain development, we performed Western blot analyses using mouse cortical lysates from different postnatal time points (Postnatal day (P) 1 through 60). Interestingly, full-length NCAN at 250KDa was only present in the lysates from P1 and P7 brains. In contrast, N and C terminal fragments were abundant across development, coinciding with the period of synapse formation and remaining high into adulthood (Figure 2B). NCAN’s developmentally timed cleavage suggests that the resulting N- and C-terminal fragments could play independent roles in the developing brain.

Previous studies showed that *Ncan* mRNA expression is highly enriched in mouse and human astrocytes compared to other brain cell types (Figures S2A and S2B). To test whether astrocytes release full-length NCAN or the N- and C-terminal fragments *in vitro*, we analyzed the expression of each in the ACM by Western blotting. We found that rat cortical astrocytes express and release NCAN full-length; however, N- and C-terminal fragments can also be detected in the ACM *in vitro* (Figure 2C).

In humans, common variations in *NCAN* are a risk factor for bipolar disorder and Schizophrenia^24,25,39^. Therefore, we decided to analyze the expression of NCAN *in vivo* in the Anterior Cingulate Cortex (ACC), an area of the brain where aberrant neuronal connectivity has been associated with the pathophysiology of SCZ and BD^40–43^. Using fragment-specific antibodies, we immunostained for NCAN in the ACC of the Aldh1L1-eGFP mice, a mouse line in which all astrocytes express GFP ^44,45^. We found that at P7, NCAN N- and C-terminal signals are abundant within astrocytes and the ECM that surrounds them (Figure 2D). These results show that astrocytes express NCAN *in vitro* and *in vivo*, and the full-length NCAN is cleaved into two distinct fragments in the brain in a developmentally regulated manner.

To understand how N- and C-terminal fragments distribute in the ECM after cleavage, we used immunofluorescence to label them at different developmental times in the ACC. We found that at P7, the majority of immunohistochemical signals for NCAN N- and C-terminal fragments overlap; however, by P30, they exhibit distinct localizations in the extracellular matrix (Figure 2E). These observations further indicate that these two fragments may conduct distinct functions in the cerebral cortex at this later stage.

NCAN is best known as one of the main components of the condensed matrix of perineuronal nets, where it binds to hyaluronic acid through its link domains^23,28^. To determine the relative distributions of N- and C-terminal NCAN fragments with respect to the PNNs, we used super-resolution stimulated emission depletion (STED) microscopy in layers 2/3 of the ACC of P30 WT mice. As previously reported for NCAN, the NCAN N-terminal is highly enriched in PNNs and colocalizes with Wisteria floribunda agglutinin (WFA), a PNN marker. In contrast, NCAN C-terminal is not enriched in the PNNs but rather is distributed in the brain parenchyma (Figure 2F). Collectively, these results show that in the developing cortex, NCAN is highly expressed by astrocytes and undergoes a cleavage. The resulting N- and C-terminal fractions differentially localize in the ECM and thus may interact with distinct subcellular structures in the cerebral cortex.

### NCAN loss impairs inhibitory synapse formation and function *in vivo*

To study if NCAN is required for inhibitory synapse formation *in vivo*, we generated a transgenic knock-out mouse line using CRISPR-Cas9. We opted to study NCAN function in a null mouse because NCAN is rapidly upregulated starting at birth and has a long half-life with an extremely slow protein turnover (90 days) ^46^. Therefore, a conditional targeting approach in astrocytes may be ineffective at depleting NCAN during synaptogenesis when using a floxed allele and a postnatal inducible-Cre driver specific to astrocytes, the Aldh1L1-CreERT2. Importantly, NCAN expression is restricted to the nervous system, and astrocytes are the primary cells expressing it^37,47–49^.

We designed the *Ncan*-null allele (*Ncan* KO) by deleting the exons 3 and 4 via a CRISPR/Cas9-mediated targeting strategy. Loss of these exons results in an mRNA transcript containing a premature STOP codon in exon 5, expected to cause nonsense-mediated decay of the mRNA (Fig 3A). Using P10 *Ncan* KO mice, we confirmed the successful deletion of *Ncan* via genomic PCR and the loss of both N and C terminal NCAN fragments in the cerebral cortex by immunofluorescence and Western blotting (Figures 3B-D).

**Figure 3:**
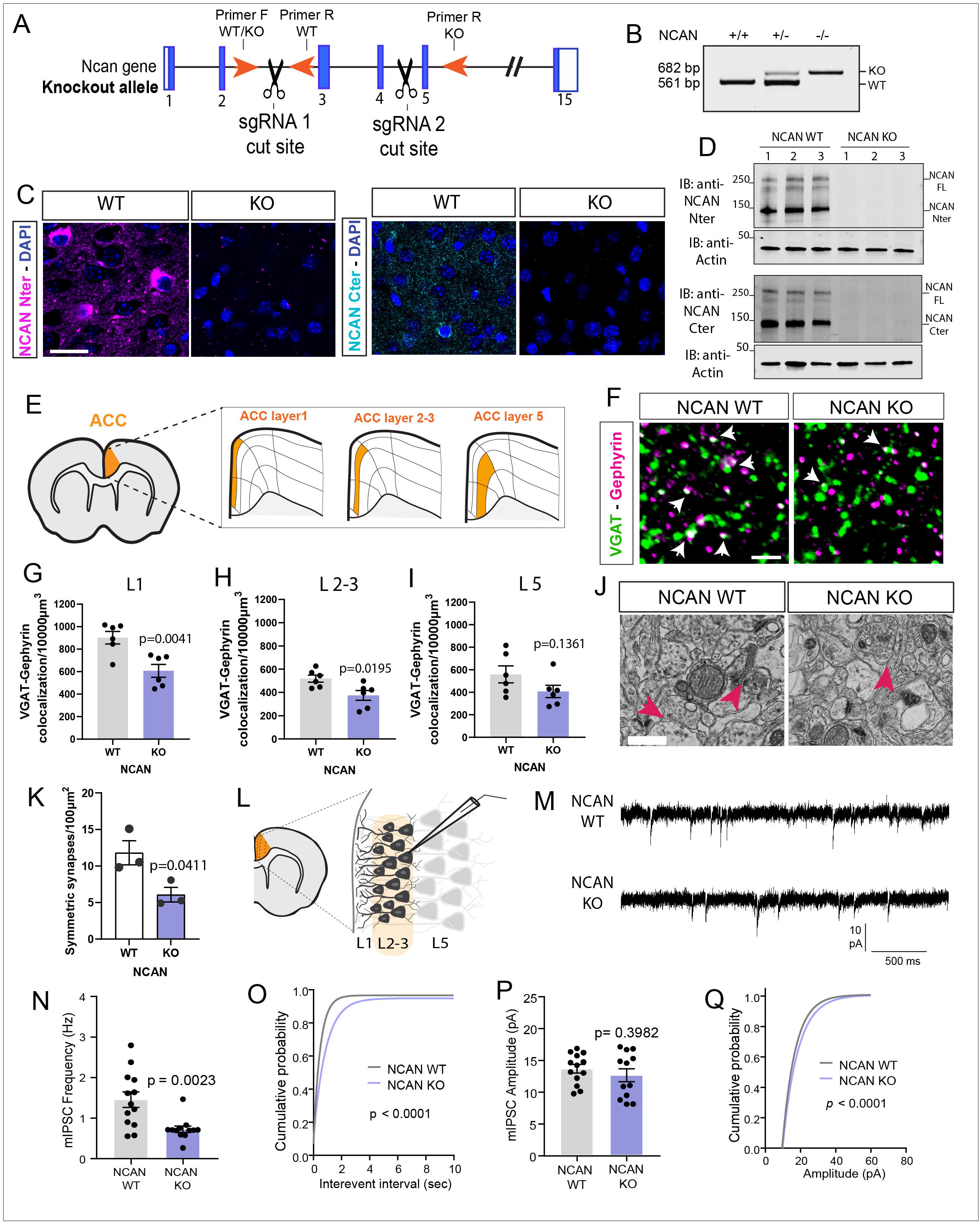
NCAN loss impairs inhibitory synapse formation and function *in vivo.* (A) Scheme of CRISPR-Cas9 strategy to delete exons 3 and 4 from the *Ncan* gene. (B) Genomic PCR from NCAN WT, Het and KO mice. (C) Representative images of NCAN N- and C-terminal in cortical sections of NCAN WT and KO mice. Scale bar: 10μm. (D) Western blot analysis of NCAN N- and C-terminal expression in cortical lysates from NCAN WT and KO mice. (E) Schematic of layer1, 2/3 and 5 in the ACC. (F) Representative images of inhibitory synapses marked with VGAT (green) and Gephyrin (magenta) in layer 2/3 of the ACC of NCAN WT and KO mice. White arrows point at inhibitory synapses. Scale bar, 10μm. (G, H and I) Quantification of the density of inhibitory synapses per image between NCAN WT and KO littermates in layer 1 (G), layer 2-3 (H) and layer 5 (I). 5 images/section, 3 sections/mouse, 6 sex-matched littermate pairs. Data points represent mouse averages. Bars are mean ± s.e.m. Unpaired two-tailed t-test. (J) Representative images of electron microscopy from the ACC from NCAN WT and KO mice. Arrowheads indicate inhibitory (symmetric) synapses. Scale Bar: 2μm. (K) Quantification of symmetric synapses in NCAN WT and KO littermates from the ACC. 10 images/mouse, 3 sex-matched littermate pairs. Data points represent mouse averages. Bars are mean ± s.e.m. Unpaired two-tailed t-test. (L) Scheme of electrophysiology recordings from layers 2-3 from the ACC. (M) mIPSC traces from L2-3 pyramidal neurons in acute ACC slices from NCAN WT and KO mice. (N and O) Quantification of frequency average (N) and cumulative probability (O) of mIPSC from NCAN WT and KO pyramidal neurons. n = 13 WT and 12 KO neurons from 4 mice per genotype. Kolmogorov-Smirnov test (D = 0.39, *p* < 0.0001). Average frequency of mIPSC in NCAN WT (1.455 ± 0.1912) and NCAN KO (0.7258 ± 0.0785) mice. Unpaired Two tailed t-test [t (23) = 3.422, *p* = 0.0023]. (P and Q) Quantification of amplitude (P) and cumulative probability (Q) from NCAN WT and KO pyramidal neurons. n = 13 WT and 12 KO neurons from 4 mice per genotype. Kolmogorov-Smirnov test (D = 0.322, *p* < 0.0001). Average amplitude of mIPSC in NCAN WT (13.7 ± 0.6636) and NCAN KO (12.66 ± 1.032) mice. Unpaired t-test [t (23) = 0.8606, *p* = 0.3982]. Data are presented as mean ± s.e.m.

Loss of NCAN did not affect cortical lamination of the ACC as we found no differences in the cortical layer thickness between P1 NCAN KO and WT mice by immunofluorescence using the cortical neuronal markers Lhx2, Ctip2, and Tbr1 (Figures S3A-F). Because NCAN has a role in inhibitory synapse formation *in vitro*, we analyzed the density of inhibitory neurons in NCAN KO mice. We found no difference in the number of GAD67-positive neurons between P30 NCAN WT and KO mice in the ACC (Figure S3G and S3H). Finally, to determine if *Ncan* deletion affected the number of different cell types in the cortex, we performed automated image segmentation based on nuclear marker labeling (see Methods for details). We found no alteration in the number or density of neurons (NeuN+), astrocytes (Sox9+/Olig2−), or oligodendrocytes (Olig2+) in the ACC of NCAN KO mice at P30 (Figures S3I and S3J). These results show that loss of NCAN does not cause gross cytoarchitectural deficits or neuron loss in the ACC.

To study NCAN’s requirement for inhibitory synapse development *in vivo*, we analyzed the density of inhibitory synapse structures, marked by the close apposition of presynaptic VGAT and postsynaptic Gephyrin, in NCAN KO mice and their littermate, sex-matched WT siblings in different layers of the ACC. We found a significant reduction in the density of inhibitory synapses (~30%) in NCAN KO mice in layers 1 and 2/3 of the ACC compared to the WT littermates with no sex-specific effect (Figure 3F-I and S3K). To confirm these histological findings at the ultrastructural level, we performed Electron Microscopy (EM) to visualize synaptic structures in the ACCs of littermate P30 WT and KO mice. To determine how NCAN loss affected inhibitory synapse numbers, we counted symmetric synapse density and compared it between genotypes. In agreement with our immunofluorescence data, the EM results indicate that NCAN KO mice present a ~50% reduction in inhibitory synapse numbers compared to WT littermates (Figure 3J and 3K). Altogether these findings show that NCAN is essential for inhibitory synapse number in the developing mouse ACC.

Our results show that NCAN controls inhibitory synaptic numbers *in vitro* and *in vivo*. Therefore, we next investigated if NCAN loss affects inhibitory synapse function as well. To do so, we performed whole-cell patch-clamp recordings of miniature inhibitory postsynaptic currents (mIPSCs) from in layers 2/3 pyramidal neurons of the ACC, using acute brain slices of P30 NCAN WT and KO mice (Figure 3L). We found a significant decrease (decreased by ~50%) in the frequency of miniature IPSCs in NCAN KO mice compared to WT littermates (Figures 3M-O), with no change in the mean amplitudes of the mIPSCs (Figures 3P and 3Q). These physiological findings are in line with the reduced numbers of inhibitory synapses we observed by immunohistochemistry and EM. Taken together, our results reveal that NCAN is required for proper inhibitory synapse formation and function *in vivo*.

### NCAN C-terminal fragment is necessary and sufficient for inducing inhibitory synapse formation *in vitro*

Because NCAN is cleaved into N and C terminal fragments during postnatal brain development and because these fragments are differentially localized within the brain parenchyma, we next interrogated the roles of NCAN’s N- and C-terminal fragments on inhibitory synapse formation. To do so, we produced recombinant proteins composed of NCAN’s protein-protein interaction domains localized to N and C terminal fragments. For these experiments, the regions of NCAN containing GAG chains were excluded. We produced two recombinant NCAN fragments, one containing the N-terminal Ig-module and Link domains (referred to as Nter-IL-domain) and the other one composed of the C-terminal EGF-like domains, Lectin-like domain, and Sushi domain (referred to as Cter-ELS-domain) (Figures 4A-C). The recombinant proteins were expressed by HEK293 cells and were purified from the conditioned media by Histidine-tag affinity purification (Figure S4A).

**Figure 4:**
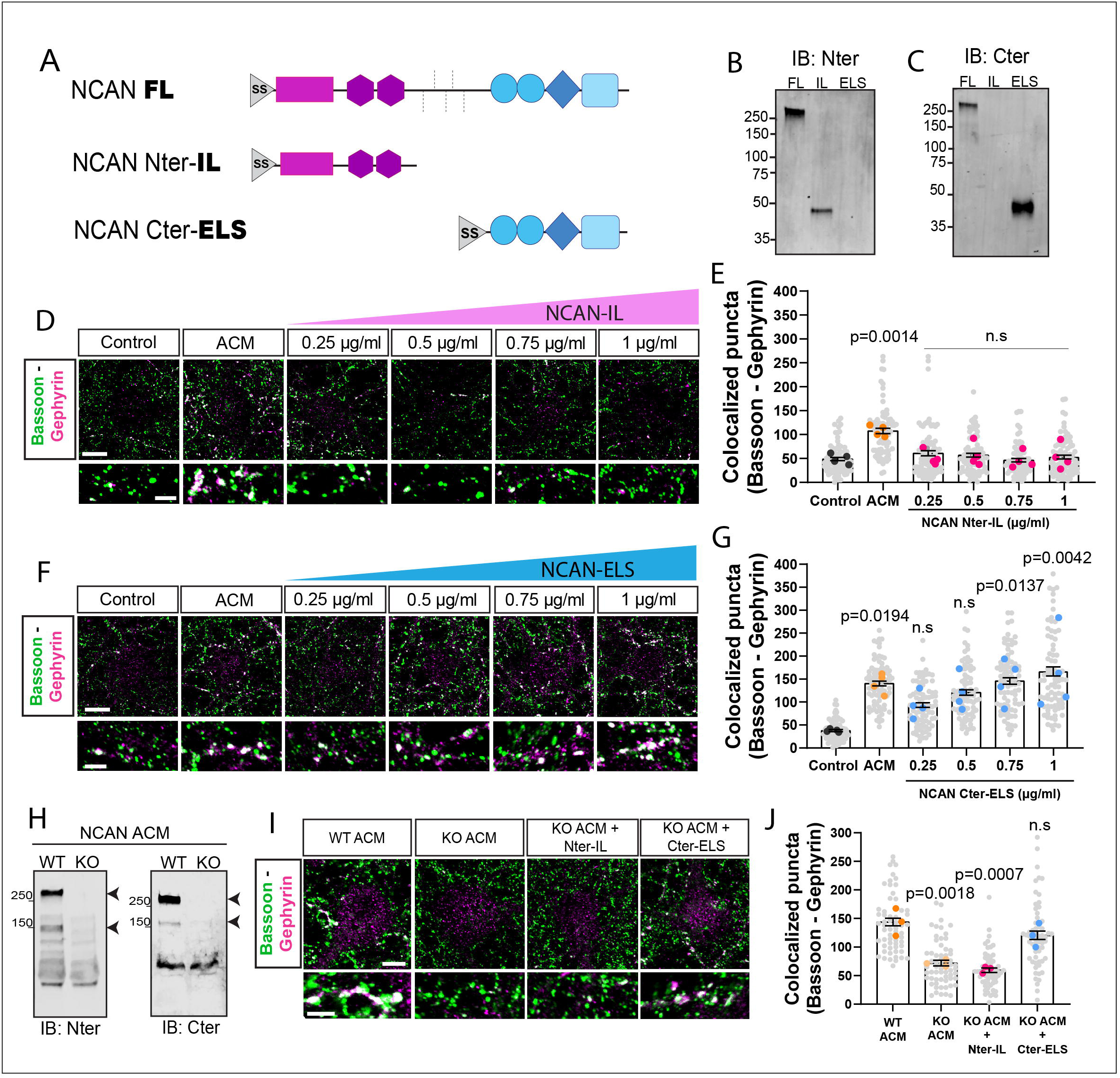
NCAN Cter but not Nter is synaptogenic *in vitro*. (A) Scheme of NCAN full-length (FL), N-terminal-IL (IL) and C-terminal-ELS (ELS) recombinant proteins. SS: secretion signal. (B) Western blot validation of NCAN-FL, IL and ELS recombinant proteins. (D) Dose-response curve of increasing concentration of NCAN-IL recombinant protein in glia-free neuronal cultures. Inhibitory synapses were marked with presynaptic marker Bassoon (green) and postsynaptic marker Gephyrin (magenta). Scale bar: 15μm. Insert scale bar: 5μm. (E) Quantification of inhibitory synapses (Bassoon and Gephyrin colocalization) with increasing concentrations of NCAN-IL recombinant protein. Data are mean ± s.e.m. n = 4 independent experiments, 20 cells/condition/experiment. One-way ANOVA, Dunnett’s post-test. (F) Dose-response curve of increasing concentration of NCAN-ELS recombinant protein in glia-free neuronal cultures. Inhibitory synapses were marked with presynaptic marker Bassoon (green) and postsynaptic marker Gephyrin (magenta). Scale bar: 15μm. Insert scale bar: 5μm. (G) Quantification of inhibitory synapses (Bassoon and Gephyrin colocalization) with increasing concentrations of NCAN-ELS recombinant protein. Data are mean ± s.e.m. n = 4 independent experiments, 20 cells/condition/experiment. One-way ANOVA, Dunnett’s post-test. (H) Western blot analysis of NCAN N- and C-terminal expression in NCAN WT and KO astrocyte-conditioned media. (I) Rescue experiment with NCAN WT ACM, KO ACM and KO ACM with NCAN IL or ELS in glia-free neuronal cultures. Inhibitory synapses were marked with presynaptic marker Bassoon and postsynaptic marker Gephyrin. Scale bar: 15μm. Insert scale bar: 5μm. (J) Quantification of inhibitory synapses (Bassoon and Gephyrin colocalization) from panel I. Data are mean ± s.e.m. n = 3 independent experiments, 20 cells/condition/experiment. One-way ANOVA, Dunnett’s post-test.

To determine whether these recombinant NCAN fragments are sufficient to induce inhibitory synapse formation in glia-free cortical cultures *in vitro*, we treated DIV 8-11 purified cortical neuron cultures with increasing concentrations of Nter-IL or Cter-ELS-domain proteins. Treatment with the N-terminal IL-domain did not increase the number of inhibitory synapses marked with Bassoon and Gephyrin compared to neurons that were cultured in growth media (Figures 4D and 4E). However, the recombinant C-terminal ELS-domain strongly induced inhibitory synapse formation at 750 ng/ml concentration, closely mimicking the effect of the ACM (Figure 4F and 4G).

We next tested whether NCAN C-terminal ELS-domain is necessary for the synaptogenic function of the mouse ACM. To do so, we tested whether ACM made from NCAN KOs retains any inhibitory synaptogenic function. First, we validated by Western blot that NCAN is present in WT mouse ACM but is absent in the ACM from NCAN KO astrocytes (Figure 4H). Second, we quantified the number of inhibitory synapses formed onto neurons in our glia-free cultures when they are exposed to WT or NCAN KO ACM. We found that NCAN KO ACM did not induce a significant increase in the number of inhibitory synapses formed onto excitatory cortical neurons when compared to the WT ACM (Figure 4I and 4J). To determine if the Cter-ELS-domain can rescue the inhibitory synaptogenic function of the NCAN KO ACM, we added it back at 750 ng/ml and analyzed inhibitory synaptic density. Reintroducing the NCAN Cter-ELS-domain rescued the inhibitory synaptic deficit of the NCAN KO ACM but adding back the Nter-IL-domain (750 ng/ml) did not (Figure 4I and 4J). Collectively, our results show that NCAN C-terminal ELS-domain is the necessary and sufficient part of NCAN that induces inhibitory synapse formation *in vitro*.

### Inhibitory synaptogenesis is impaired in NCAN-ΔELS mutant mice

To analyze the functions of the NCAN C-terminal ELS-domain *in vivo*, we designed a second mouse line, NCAN-ΔELS mutant using the CRISPR-Cas9 system. We targeted exons 9 through 14 of the *Ncan* gene to remove the EGF-like, Lectin-like, and Sushi domains, leaving the rest of the gene unaltered (Figures 5A and 5B). We performed genomic PCR, immunofluorescence, and Western blot to validate this new mutant mouse line (Figure 5C-E). Indeed, NCAN-ΔELS mice express an intact NCAN N-terminal domain but lack the expression of the ELS-domains at the C-terminal (Figures S5A, 5C, and 5D). By immunofluorescence, we found that the localization of the N-terminal fragment at the PNNs is not affected in the ΔELS mutant mice (Figure 5E). These results show that targeting exons 9 through 14 in *Ncan* results in a mutant mouse that expresses an intact N-terminal fragment but lacks the synaptogenic Cter-ELS domain.

**Figure 5:**
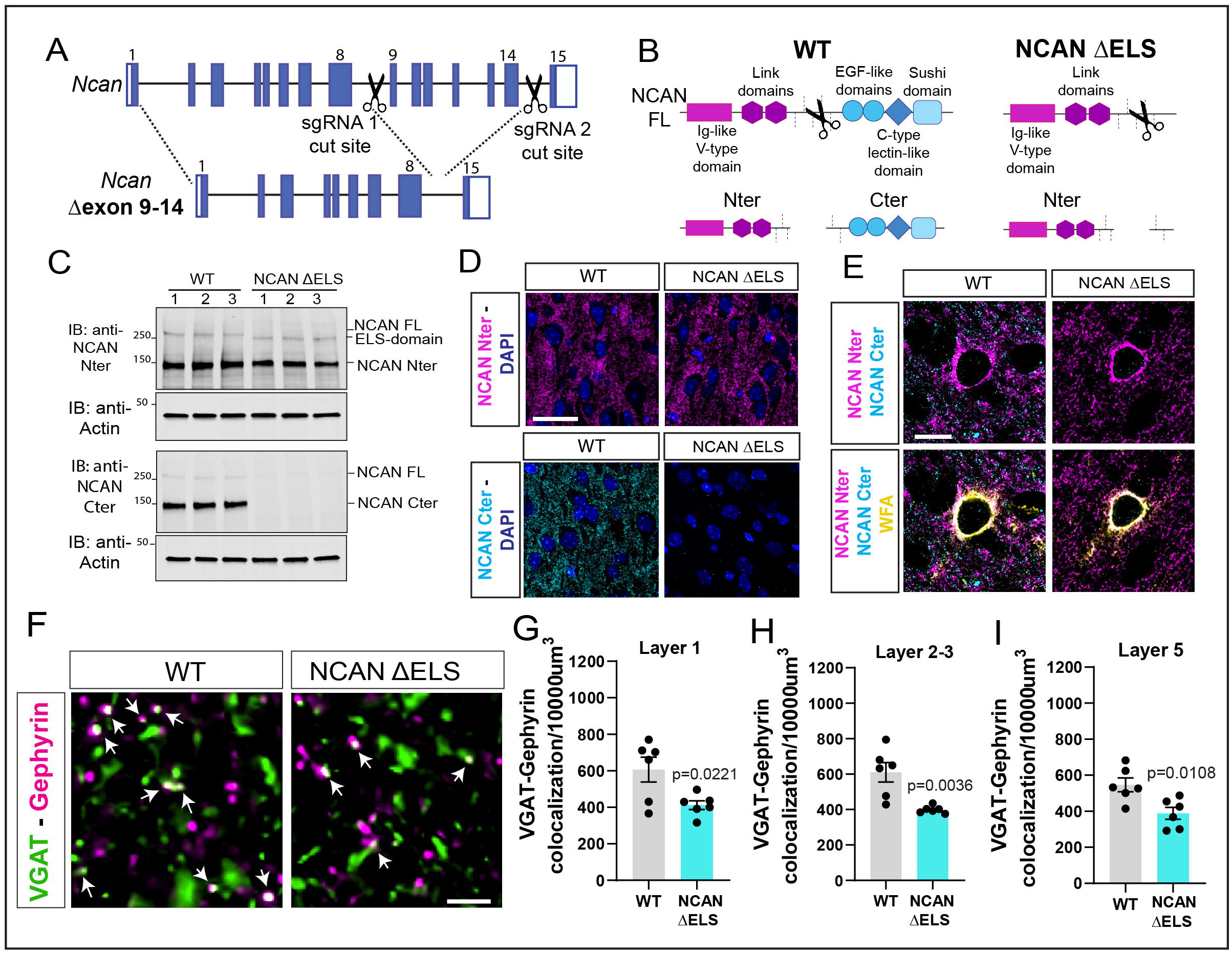
Inhibitory synaptogenesis is impaired in NCAN ΔELS mutant mice. (A) Scheme of CRISPR-Cas9 strategy to delete exons 9 to 14 from the *Ncan* gene. (B) Scheme of NCAN WT protein and NCAN ΔELS mutant. (C) Western blot analysis of NCAN N- and C-terminal expression in cortical lysates from NCAN WT and ΔELS mutant mice. (D) Representative images of NCAN N- and C-terminal in cortical sections of NCAN WT and ΔELS mutant mice. Scale bar: 100μm. (E) Representative images of NCAN N- and C-terminal distribution at perineuronal nets marked with WFA (yellow) in cortical sections of NCAN WT and ΔELS mutant mice. Scale bar: 30μm. (F) Representative images of inhibitory synapses marked with VGAT (green) and Gephyrin (magenta) in layer 2/3 of the ACC of NCAN WT and ΔELS mutant mice. Scale bar, 10μm. (G, H and I) Quantification of the density of inhibitory synapses per image between NCAN WT and ΔELS mutant littermates in layer 1 (G), layer 2-3 (H) and layer 5 (I). 5 images/section, 3 sections/mouse, 6 sex-matched littermate pairs. Data points represent mouse averages. Bars are mean ± s.e.m. Unpaired two-tailed t-test.

We further validated this new mutant mouse model by determining if loss of the Cter-ELS-domain alters gross brain morphology. We found no differences in the numbers of inhibitory neurons marked with GAD67 in the ACCs of P30 NCAN-ΔELS mutant mice and their WT littermates (Figures S5B and S5C). Similarly, we found no changes in the densities of neurons (NeuN+), astrocytes (Sox9+/Olig2−), or oligodendrocytes (Olig2+) in the ACC at P30 between littermate NCAN-ΔELS mutant mice and WT mice (Figures S5D and S5E). These results show that the deletion of the NCAN ELS domain does not cause changes in the number of neural cell types.

To determine whether NCAN Cter-ELS-domain is required for proper inhibitory synaptogenesis in the developing brain, we quantified the inhibitory synapse density in the ACC of littermate WT and NCAN-ΔELS mice at P30 (Figure 5F). In NCAN-ΔELS mice, we found a marked decrease (~30%) in the density of inhibitory synapses in layers 1, 2/3, and 5 (Figures 5G-I), consistent with what we observed in the NCAN KO mice. These results show that the NCAN ELS domain is required for inhibitory synapse formation *in vivo*.

### NCAN ELS domain interacts with excitatory and inhibitory synaptic proteins

NCAN ELS-domain contains protein-protein interaction motifs commonly associated with cell adhesion ^50–52^. We hypothesized that the NCAN C-terminal fragment could control inhibitory synapse formation and function by interacting with neuronal proteins localized to synapses. Using STED microscopy, we found that the NCAN C-terminal fragment colocalizes with 75% of inhibitory synapses (marked by colocalization of VGAT-Gephyrin puncta) and 40% of excitatory synapses (marked by colocalization of VGluT1-Homer1 puncta) (Figure 6A-D). On the contrary, the NCAN N-terminal fragment is specifically enriched at PNNs (Figure 2F) and is less abundant at inhibitory synapses (Figure S6A and S6B).

**Figure 6:**
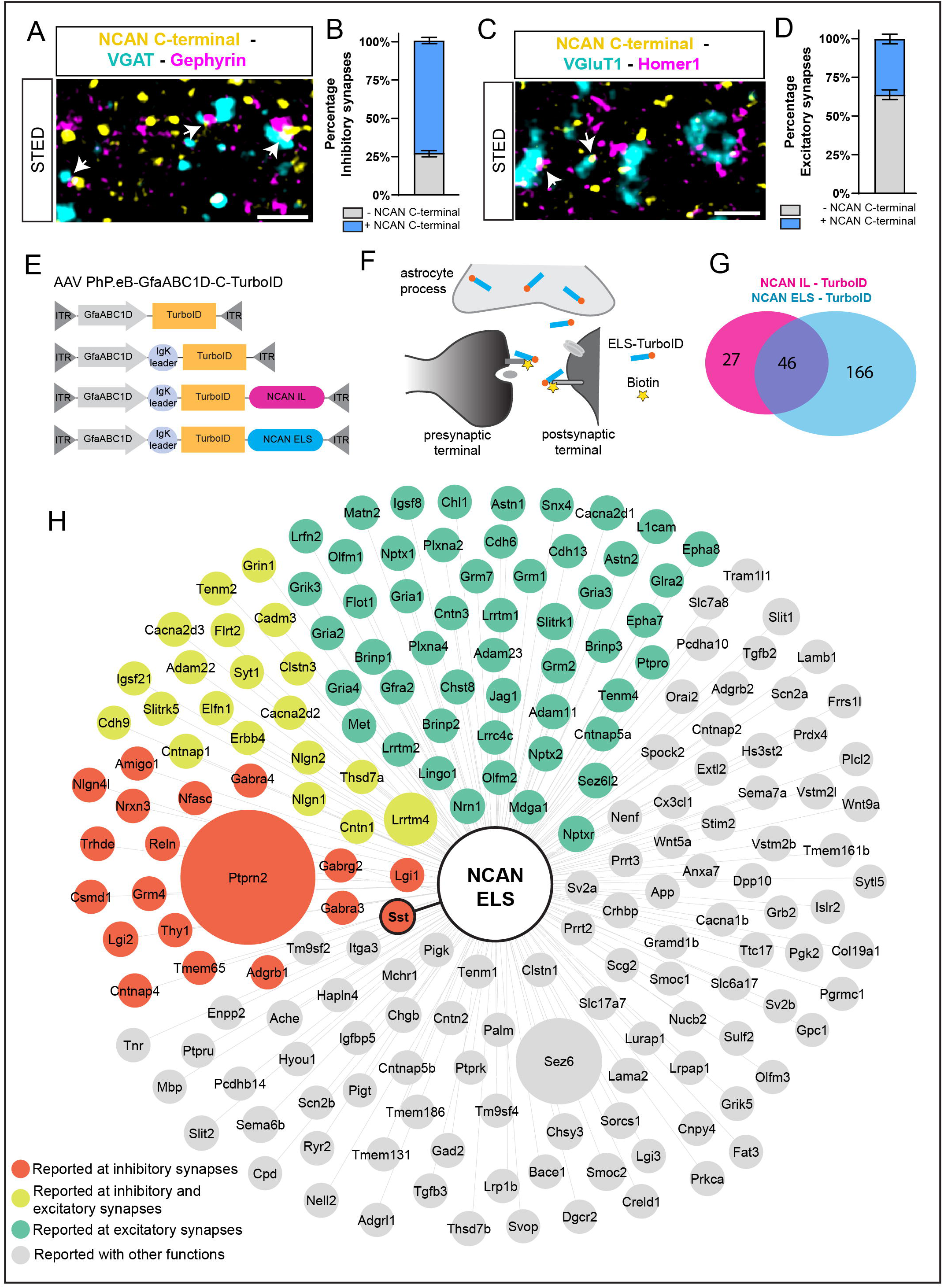
NCAN ELS domain interactome contains proteins enriched at excitatory and inhibitory synapses. (A) Three-color STED image of NCAN C-terminal together with VGAT and Gephyrin in P30 WT mice, layer 2-3 of the ACC. Scale bar: 1μm. (B) Percentage of NCAN C-terminal colocalizing with inhibitory synapses. 3 sections/mouse, 3 WT CD1 mice. Bars are mean ± s.e.m. (C) Three-color STED image of NCAN C-terminal together with VGluT1 and Homer1 in P30 WT mice, layer 2-3 of the ACC. Scale bar: 1μm. (D) Percentage of NCAN C-terminal colocalizing with excitatory synapses. 3 sections/mouse, 3 WT CD1 mice. Bars are mean ± s.e.m. (E) Representation of TurboID constructs. IgK leader: secretion signal. (F) Schematic of the secreted TurboID approach. (G) Venn diagram of NCAN IL and ELS extracellular proteome. (H) NCAN ELS-TurboID secreted and membrane receptor enriched proteins.

To identify the extracellular binding partners of the NCAN ELS domain, we developed an *in vivo* chemicogenetic approach. Proximity biotin labeling by TurboID enzyme has been shown to be a fast, reliable, and non-toxic *in vivo* approach for labeling endogenous interacting proteins ^53–59^. However, so far, this approach has been mostly utilized to identify the interactome of intracellular proteins^54,58^. To apply this strategy to NCAN, we virally expressed in astrocytes (GfaABC1D promoter) a secreted version of the TurboID enzyme fused to the NCAN Cter-ELS or the NCAN Nter-IL domains (Figure 6E and 6F). To control for unspecific interaction partners of these fragments, we expressed cytosolic TurboID. On the other hand, to eliminate the interactions of NCAN domains within the secretory organelles of astrocytes, we also used a secreted Turbo ID containing the same secretion sequence (IgK leader sequence) in astrocytes (Figure 6E and 6F). These viruses were tested *in vitro* for their ability to be expressed in astrocytes and secrete the NCAN fragments to the ACM and biotinylate proteins (Figure S6C and S6D).

To identify NCAN interaction partners, each virus was intracranially injected into the cerebral cortices of P1 WT mice. At P18, mice received biotin injections for 3 days until P21 (Figure S6E) at which point the cortices were collected. This experimental design allowed for the broad expression of the TurboID constructs in astrocytes resulting in abundant biotinylation (Figure S6F). Biotinylated proteins were purified from cortical lysates for proteomic analysis (Figure S6G).

Biotinylated proteins were analyzed by quantitative high-resolution liquid chromatography– tandem mass spectrometry (LC–MS)^54,56^. From this analysis, 4726 unique proteins were identified (Table S1). Compared with secreted IgK-TurboID and Cytosolic-TurboID controls, we found 27 and 166 unique membrane and secreted proteins significantly enriched (> 1.5-fold) in NCAN Nter-IL-TurboID and NCAN Cter-ELS-TurboID fractions, respectively (Figure 6G). Moreover, we found 46 shared proteins between Nter-IL and Cter-ELS (Figure 6G and Table S2). We filtered these proteomic lists for known NCAN Nter-IL and Cter-ELS interactors and found that both NCAN Nter-IL-TurboID and NCAN Cter-ELS-TurboID interacting proteins contain those present at PNNs (Figure S6H). Next, we analyzed if NCAN Cter-ELS interactome contains known synaptic proteins. To do that, we visualized the 166 unique NCAN Cter-ELS interactors together with proteins from the 46 shared NCAN Nter-IL and Cter-ELS samples that were at least 1.5-fold-enriched in their interaction with the Cter-ELS domain (Figure 6H, Table S2 and S3). We found this final proteomic list includes known synaptic proteins from excitatory synapses: cell adhesion molecules Lrrtm1/2/4; AMPA receptors (*Gria1*, *Gria2* and *Gria3*); and proteins present at inhibitory synapses, such as the leucine-rich secreted protein Lgi1/2 and GABA receptors (G*abra3* and 4) (Figure 6H). Interestingly, NCAN ELS interacts with the neuropeptide somatostatin, which is expressed by SST+ GABAergic interneurons in the neocortex^60^. These results revealed that in the extracellular matrix NCAN IL and ELS interactome diverge, and NCAN ELS interacts with multiple synaptic proteins. Because NCAN ELS is in proximity with SST, we decided to investigate if NCAN C-terminal fragment controls the formation of distinct inhibitory synaptic circuits.

### NCAN C-terminal controls SST synapse formation and function

To study if NCAN C-terminal differentially localizes to SST+ or PV+ synapses, we used STED super-resolution microscopy. We labeled SST+ synapses with antibodies against presynaptic marker SST and postsynaptic marker Gephyrin. To visualize PV+ synapses, we used presynaptic maker Synaptotagmin2 (Syt2), which labels axo-somatic PV+ synapses from basket cells onto neuronal bodies^6^, together with postsynaptic Gephyrin (Figure 7A). We counted the number of SST+ and Syt2+ synapses colocalizing with NCAN C-terminal fragments in P30 WT mice in layer 2/3 of the ACC. We found that 50% of SST+ synapses colocalize with the NCAN C-terminal domain; however, only 25% of Syt2+ synapses colocalize with this fragment (Figure 7B-E), indicating that the C-terminal domain is enriched at SST+ synaptic contacts.

**Figure 7:**
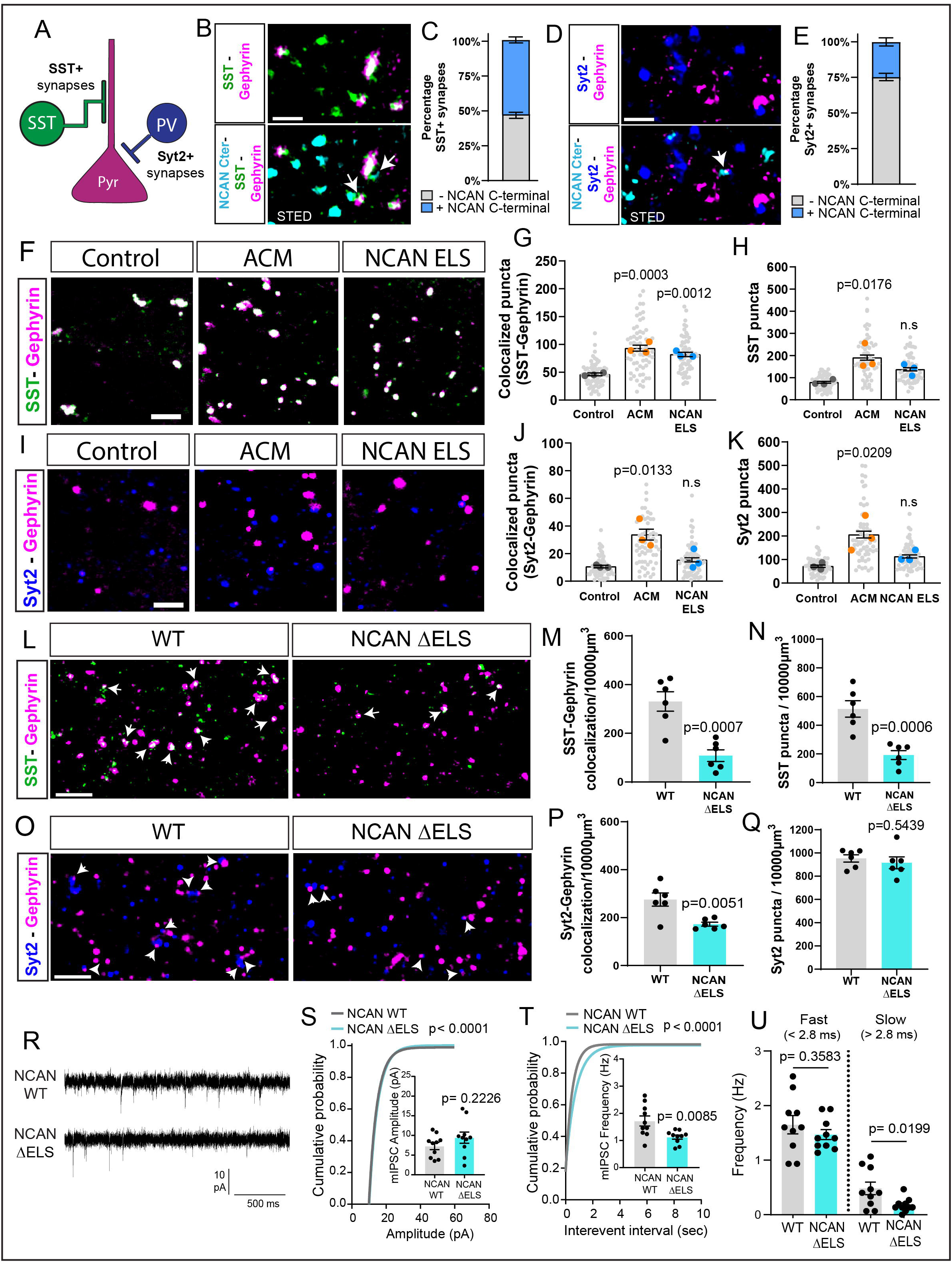
NCAN C-terminal fragment controls SST synaptogenesis. (A) Scheme of somatostatin (SST) and PV (Syt2) targets on pyramidal glutamatergic neurons. (B) Three-color STED image of NCAN C-terminal together with SST and Gephyrin in P30 WT mice, layer 2-3 of the ACC. White arrows indicate synapses with NCAN Cter. Scale bar: 1μm. (C) Percentage of distribution of NCAN C-terminal colocalizing with SST+ synapses. 3 sections/mouse, 3 WT CD1 mice. Bars are mean ± s.e.m. (D) Three-color STED image of NCAN C-terminal together with Syt2 and Gephyrin in P30 WT mice, layer 2-3 of the ACC. White arrows indicate synapses with NCAN Cter. Scale bar: 1μm. (E) Percentage of distribution of NCAN C-terminal colocalizing with Syt2+ synapses. 3 sections/mouse, 3 WT CD1 mice. Bars are mean ± s.e.m. (F) Representative images of SST+ synapses in glia-free neuronal cultures treated with control, ACM or NCAN ELS recombinant protein. SST+ inhibitory synapses were marked with presynaptic marker SST (green) and postsynaptic marker Gephyrin (magenta). Scale bar: 5μm. (G) Quantification of SST+ inhibitory synapses (SST and Gephyrin colocalization) from F. Data are mean ± s.e.m. n = 3 independent experiments, 20 cells/condition/experiment. One-way ANOVA, Dunnett’s post-test. (H) Quantification of SST puncta from F. Data are mean ± s.e.m. n = 3 independent experiments, 20 cells/condition/experiment. One-way ANOVA, Dunnett’s post-test. (I) Representative images of Syt2+ synapses (PV) in glia-free neuronal cultures treated with control, ACM or NCAN ELS recombinant protein. Syt2+ inhibitory synapses were marked with presynaptic marker Syt2 (green) and postsynaptic marker Gephyrin (magenta). Scale bar: 5μm. (J) Quantification of Syt2+ inhibitory synapses (Syt2 and Gephyrin colocalization) from I. Data are mean ± s.e.m. n = 3 independent experiments, 20 cells/condition/experiment. One-way ANOVA, Dunnett’s post-test. (K) Quantification of Syt2 puncta from F. Data are mean ± s.e.m. n = 3 independent experiments, 20 cells/condition/experiment. One-way ANOVA, Dunnett’s post-test. (L) Representative images of SST+ inhibitory synapses marked with SST (green) and Gephyrin (magenta) in layer 2-3 of the ACC of NCAN WT and ΔELS mutant mice. White arrows indicate SST+ synapses. Scale bar, 10μm. (M) Quantification of the density of SST+ inhibitory synapses in NCAN WT and ΔELS mutant littermates in the ACC layer 2-3. 5 images/section, 3 sections/mouse, 6 sex-matched littermate pairs. Data points represent mouse averages. Bars are mean ± s.e.m. Unpaired two-tailed t-test. (N) Quantification of SST puncta in NCAN WT and ΔELS mutant littermates in the ACC layer 2-3. 5 images/section, 3 sections/mouse, 6 sex-matched littermate pairs. Data points represent mouse averages. Bars are mean ± s.e.m. Unpaired two-tailed t-test. (O) Representative images of PV (Syt2+) inhibitory synapses marked with Syt2 (blue) and Gephyrin (magenta) in layer 2/3 of the ACC of NCAN WT and ΔELS mutant mice. White arrows indicate Syt2+ synapses. Scale bar, 10μm. (P) Quantification of the density of Syt2+ inhibitory synapses in NCAN WT and ΔELS mutant littermates in the ACC layer 2-3. 5 images/section, 3 sections/mouse, 6 sex-matched littermate pairs. Data points represent mouse averages. Bars are mean ± s.e.m. Unpaired two-tailed t-test. (Q) Quantification of Syt2 puncta in NCAN WT and ΔELS mutant littermates in the ACC layer 2-3. 5 images/section, 3 sections/mouse, 6 sex-matched littermate pairs. Data points represent mouse averages. Bars are mean ± s.e.m. Unpaired two-tailed t-test. (R) mIPSC traces from L2-3 pyramidal neurons in acute ACC slices from NCAN WT and ΔELS mutant mice. (S) Quantification of amplitude and cumulative probability from NCAN WT and ΔELS mutant mice pyramidal neurons. n = 10 WT and 10 ΔELS neurons from 3 mice per genotype. Kolmogorov-Smirnov test (D = 0.18, *p* < 0.0001). Average amplitude of mIPSC in NCAN WT (7.322 ± 0.9048) and ΔELS mutant (9.44 ± 1.411) mice. Unpaired t-test [t (18) = 1.263, *p* = 0.2226]. (T) Quantification of frequency average and cumulative probability of mIPSC from NCAN WT and ΔELS mutant mice pyramidal neurons. n = 10 WT and 10 ΔELS neurons from 3 mice per genotype. Kolmogorov-Smirnov test (D = 0.266, *p* < 0.0001). Average frequency of mIPSC in NCAN WT (1.718 ± 0.1859) and ΔELS mutant (1.125 ± 0.07633) mice. Unpaired Two tailed t-test [t (18) = 2.952, *p* = 0.0085]. (U) Frequency of fast somatic, <2.8ms) and slow dendritic (>2.8ms) events in L2-3 pyramidal from NCAN WT and ΔELS mutant mice. n = 10 WT and 10 KO neurons from 3 mice per genotype. Average frequency of fast somatic events in NCAN WT (1.647 ± 0.1678) and ΔELS mutant (1.467 ± 0.09108) mice. Unpaired Two tailed t-test [t (18) = 0.9428, *p* = 0.3583]. Average frequency of slow dendritic events in NCAN WT (0.48 ± 0.1132) and ΔELS mutant (0.1733 ± 0.04) mice. Unpaired Two tailed t-test [t (18) = 2.544, *p* = 0.0199]. Data are mean ± s.e.m.

To study if NCAN ELS induces the formation of different inhibitory synapse subtypes, we cultured glia-free cortical neurons with ACM or recombinant NCAN ELS protein. We quantified the number of SST+ or Syt2+ synapses as the co-localization of SST/Gephyrin or Syt2/Gephyrin (Figure 7F and 7I). We found that ACM treatment increases the numbers of both SST+ and PV Syt2+ synapses *in vitro*. However, NCAN ELS strongly increased SST+ synapses without affecting Syt2+ synapse numbers (Figures 7F-G and 7I-J). NCAN ELS domain had no effect on the numbers of Syt2 or SST+ presynaptic puncta *in vitro*. However, the ACM treatment caused a significant increase in SST and Syt2 puncta (Figure 7H and 7K), indicating that ACM may contain additional factors that control inhibitory synapse formation.

Next, we analyzed if the ablation of the NCAN ELS-domain affects the numbers of SST+ or PV+ synapses *in vivo*. We quantified the synaptic density of these two types of inhibitory synapses in layer 2/3 of the ACC in P30 NCAN ΔELS mice and their WT littermates (Figure 7L and 7O). We found that the densities of both Syt2+ and SST+ synapses were reduced compared to the control littermates; however, the reduction in SST+ synapses was far more pronounced (Figure 7L-M and 7O-P). Intriguingly, in the mutant mice, the density of SST puncta, but not Syt2, was also significantly reduced compared to the littermate WTs (Figure 7N and 7Q), indicating that SST+ synaptic terminals are susceptible to NCAN ELS deletion. Together, these results show that NCAN ELS domain is crucial for the development of SST+ and to a lesser extent the Syt2+ PV synapses in the mouse cortex.

Finally, we investigated how the loss of ELS-domain impacts inhibitory synaptic function. We recorded mIPSCs from layer 2/3 pyramidal neurons in the ACC in P30 NCAN ΔELS mutant mice or control WT littermates (Figure 7R). Similar to what we observed in NCAN global KOs, we found no change in the mIPSC amplitude (Figures 7S and S7B). However, the frequency of the mIPSCs in the NCAN ΔELS mutant mice was significantly reduced (~35%) compared to WT littermates (Figure 7T). *In vivo*, SST+ synapses are formed mostly on the distal dendrites of pyramidal neurons, while PV+ synapses are formed mainly on the neuronal soma (perisomatic) ^61,62^. We analyzed the recording data to determine if the reduction in mIPSC frequency in the mutant mice is due to the loss of perisomatic or distal dendritic synaptic events. To do so, we took advantage of the rise time difference between perisomatic (fast) and dendritic (slow) inhibition and analyzed the rise time of mIPSCs in WT and NCAN-ΔELS mutant mice ^63,64^. We observed a decrease (~28%) in the 10-90% rise times of the mIPSCs in NCAN-ΔELS mutant mice (Figure S7A). We then classified fast mIPSCs (perisomatic events: < 2.8 ms) and slow mIPSCs (distal dendritic events: > 2.8 ms). We did not see a change in the amplitudes of fast or slow mIPSCs (Figure S7B) but found a significant decrease (decreased by ~64%) in the frequency of slow mIPSCs in NCAN-ΔELS mutant mice (Figure 7U). These results strongly suggest that the NCAN ELS domain controls the formation and function of dendrite-targeting inhibitory synapses, which are often formed by the SST+ interneuron contacts. Altogether our findings reveal a mechanism through which astrocytes control circuit-specific inhibitory synapse formation in the developing cortex.

## DISCUSSION

### Astrocytes control inhibitory synaptogenesis

The balance between excitation and inhibition in the brain must be delicately coordinated by cellular and molecular mechanisms that distinctly control excitatory versus inhibitory synaptogenesis^1,65,66^. Astrocytes powerfully influence the formation and maturation of both types of synapses through secreted synaptogenic factors and cell adhesion proteins. To date, several cell adhesion molecules and astrocyte-secreted factors that induce excitatory synapse formation have been identified ^10–13,17,19,45,67,68^. Recently it has been shown that astrocytic-neuron contacts through adhesion protein NrCAM regulate inhibitory synapse formation ^56^. However, the identity of the astrocyte-secreted signals and the molecular mechanisms through which they control inhibition in the CNS are less understood. Here we show that Neurocan, an astrocyte-enriched chondroitin sulfate proteoglycan, controls the formation and function of inhibitory synapses. NCAN expression in astrocytes peaks in the first postnatal week, a period that correlates with astrocyte maturation and the beginning of synapse formation ^37,49^. Once in the extracellular matrix, NCAN is cleaved into two fragments^28^. We revealed that the resulting N-terminal and C-terminal fragments segregate in the neuropil, indicating possible unique functions for each fragment. Our results show that NCAN C-terminal induces inhibitory synaptogenesis independently of the N-terminal fragment and the GAG-chain interactions. These results expose a new mechanism through which astrocytes control inhibitory synapse formation by secreting a synaptogenic protein that undergoes cleavage right before the peak of synaptogenesis. Moreover, our results indicate that loss of the synaptogenic domain in NCAN C-terminal fragment impairs inhibitory synaptic function, highlighting the importance of astrocyte-secreted molecules in the control of excitatory/inhibitory synaptic balance and brain connectivity.

### NCAN C-terminal processing and function

NCAN full-length protein binds hyaluronic acid through the link domains located in the N-terminal part, which is stabilized by HAPLN2^23,69^. Our super-resolution images from PNNs show that while the N-terminal fragment is highly enriched in these structures, the C-terminal domain does not show preferential enrichment, and it is equally distributed within the brain parenchyma. When present in PNNs, NCAN C-terminal associates with Tenascin-R^70^. Quadruple knockout mice deficient for Brevican, Neurocan, Tenascin-C and Tenascin-R display altered inhibitory/excitatory synaptic number, impaired PNNs structure, and reduced PV inhibitory neuron survival^71,72^, highlighting the importance of extracellular matrix proteins in the development of neuronal networks. However, in these studies, the specific contributions of each component to the final phenotypes were hard to dissect. Our results indicate that NCAN-deficient mice and ΔELS mutant mice do not present altered inhibitory neuron numbers. Moreover, the localization of NCAN N-terminal at PNNs is not affected by the deletion of the ELS fragment *in vivo*.

Our *in vivo* chemicogenetic protein labeling approach reveals that NCAN C-terminal interacts with several receptors and proteins known to be at inhibitory synapses, for example, GABA receptors and the neuropeptide SST, indicating that the C-terminal fragment also functions outside PNNs at the synaptic level. Is NCAN cleavage required for C-terminal synaptogenic effect? It is thought that NCAN processing requires proteolytic cleavage between amino acids 638 and 639^28^. However, previous studies using serine and cysteine protease inhibitors and metalloproteinase inhibitors on astrocyte cultures could not detect changes in NCAN processing^28^. In our study, we show that although at P7 by immunofluorescence NCAN N-terminal and C-terminal colocalize in the extracellular matrix, the Western blot analysis revealed that a considerable fraction is already cleaved at this time point. Therefore, NCAN undergoes a developmentally timed cleavage that occurs once it is secreted and correlates with the synaptogenic period. The identification of the molecular and cellular mechanism triggering NCAN processing is an important topic for future study.

### NCAN C-terminal synaptogenic function

In the cerebral cortex, the heterogeneity of GABAergic synapses arises from the different interneuron subtypes that form them^2,5,6,73^. The formation of inhibitory synapses relies in part on non-overlapping molecular programs between distinct interneuron subclasses. For example, LGI2, FGF13 and CBLN4 regulate the development of PV+ basket, chandelier synapses, and SST+ synapses, respectively ^6^. Can astrocytes differentially coordinate the formation of distinct GABAergic synaptic contacts? Our results reveal a mechanism by which astrocytes control SST+ inhibitory synaptogenesis and function. We show that while astrocyte-conditioned media induce both SST+ and PV+ synapses in glia-free neuronal cultures, NCAN C-terminal fragment specifically promotes the formation of SST+ synapses. However, *in vivo*, lack of ELS domain expression results in a severe deficit in SST+ and significant reduction in PV+ basket synapse density. Combined, our *in vitro* and *in vivo* results bring fort the question if NCAN C-terminal effect on PV synapses is a direct effect or an indirect consequence of reduced SST+ synaptic numbers. In fact, PV+ cell synapses maturate later in development than SST+ ones, and it requires a feed-forward circuit with SST+ neurons. Disruption of the SST interneuron network in the cortex results in impaired PV synaptic connectivity^74,75^. Other types of interneurons, like VIP+, Reelin+, or NPY+ neurons, were not tested in our analysis. Although those neurons comprise a smaller proportion of interneurons in the cortex, they form critical neuronal networks in different cortical areas and layers^73,76,77^. Therefore, future analysis should address how astrocyte-secreted molecules, together with interneuron subtype distribution, affect GABAergic synaptic diversity within distinct areas of the brain.

### Role of Neurocan on neurological disorders and brain injury

The extracellular matrix is an important regulator of the synaptic balance in the brain^47^. Once it is formed, the components and structure of the mature ECM remain stable. However, in many pathophysiological conditions and neurological disorders, the integrity and composition of the ECM are altered^78,79^. For example, previous studies in models of amyotrophic lateral sclerosis (ALS) have shown that the affected motor neurons are surrounded by accumulating levels of Neurocan and Versican, which form non-permissive microenvironments for regeneration^80^. Moreover, the expression level of NCAN C-terminal domain in the cerebrospinal fluid of patients with ALS is significantly increased compared to control patients^81^. EGF and TGFβ signaling are known to be persistently activated in ALS and are two cytokines that induce NCAN cleavage^28,82^. Therefore, targeting EGF and TGFβ pathways in ALS could serve as a therapeutic approach to ameliorate NCAN’s effect on motor neurons.

A hallmark of many neurodevelopmental and neurodegenerative disorders is an excitatory/inhibitory synaptic imbalance^65,83,84^. This is the case in Rett syndrome, in which inhibitory synaptic number and synaptic function are altered^84,85^. Proteomic analysis of astrocyte-conditioned media from a Rett mouse model (*Mecp2* knockout) showed a significant increase in NCAN expression levels compared to the control^86^. Moreover, mutations in *NCAN* are associated with impaired memory, and visuospatial skills in humans and are a risk factor for Bipolar Disorder, Schizophrenia, and major depressive disorder^24,26,39^. Interestingly, the expression of somatostatin mRNA in the dorsolateral prefrontal cortex of subjects with schizophrenia is reduced compared to control patients^87–89^. Therefore, the changes we analyzed in SST synaptic number and inhibitory synaptic function in NCAN ΔELS mutant mice could underlie the synaptic imbalance and behavioral impartments seen in SCZ in humans. In conclusion, we have identified the astrocyte-secreted NCAN C-terminal fragment as a novel inhibitory synaptogenic protein controlling SST+ synapses and inhibitory synaptic function. It serves as a promising therapeutic target with potential implications in the pathogenesis of neurological disorders and brain injury.

## FIGURE LEGENDS

**Figure S1:**
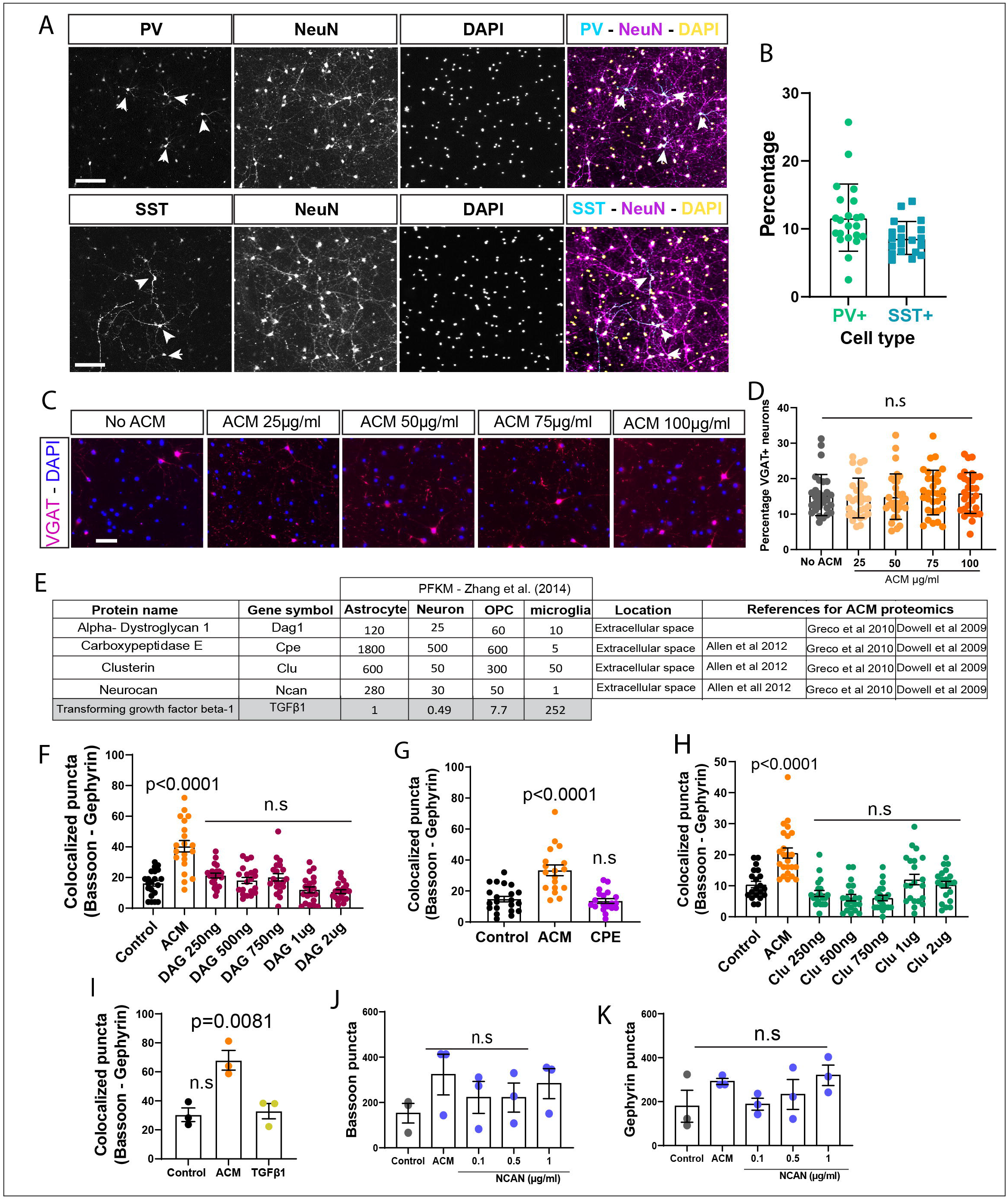
An *in vitro* screen to identify astrocyte-secreted proteins that induce inhibitory synapse formation. Related to Figure 1. (A) Representative images of PV and SST inhibitory neurons in glia-free neuronal cultures marked with neuronal marker NeuN together with SST or PV marker. Arrowheads indicate PV+ or SST+ neurons, respectively. Scale bar: 200μm. (B) Quantification of percentage of PV and SST neurons in glia-free neuronal cultures. n:1, data are mean ± s.e.m, 20 cells/condition. (C) Representative images of inhibitory neurons in glia-free neuronal cultures marked with VGAT at different concentrations of ACM. Scale bar: 100 μm. (D) Quantification of VGAT+ neurons in glia-free neuronal cultures with increasing concentration of ACM. n:1, data are mean ± s.e.m, 20 cells/condition. One-way ANOVA, Dunnett’s post-test. (E) Chart of ACM proteins selected for inhibitory synapse analysis. (F) Quantification of inhibitory synapses (Bassoon and Gephyrin colocalization) with increasing concentrations of DAG1 (ug/ml) recombinant protein in glia-free neuronal cultures. n:1, data are mean ± s.e.m, 20 cells/condition. One-way ANOVA, Dunnett’s post-test. (G) Quantification of inhibitory synapses (Bassoon and Gephyrin colocalization) with CPE (1ug/ml) recombinant protein in glia-free neuronal cultures. n:1, data are mean ± s.e.m, 20 cells/condition. One-way ANOVA, Dunnett’s post-test. (H) Quantification of inhibitory synapses (Bassoon and Gephyrin colocalization) with increasing concentrations of Clu (ug/ml) recombinant protein in glia-free neuronal cultures. n:1, data are mean ± s.e.m, 20 cells/condition. One-way ANOVA, Dunnett’s post-test. (I) Quantification of inhibitory synapses (bassoon and gephyrin colocalization) with TGFβ1 recombinant protein (10ng/ml) in glia-free neuronal cultures. Data are mean ± s.e.m. n = 3 independent experiments, 20 cells/condition/experiment. One-way ANOVA, Dunnett’s post-test. (J) Quantification of Bassoon puncta in glia-free neuronal cultures exposed to increasing concentrations of NCAN recombinant protein. Data are mean ± s.e.m. n = 3 independent experiments, 20 cells/condition/experiment. One-way ANOVA, Dunnett’s post-test. (K) Quantification of Gephyrin puncta in glia-free neuronal cultures exposed to increasing concentrations of NCAN recombinant protein. Data are mean ± s.e.m. n = 3 independent experiments, 20 cells/condition/experiment. One-way ANOVA, Dunnett’s post-test.

**Figure S2:**
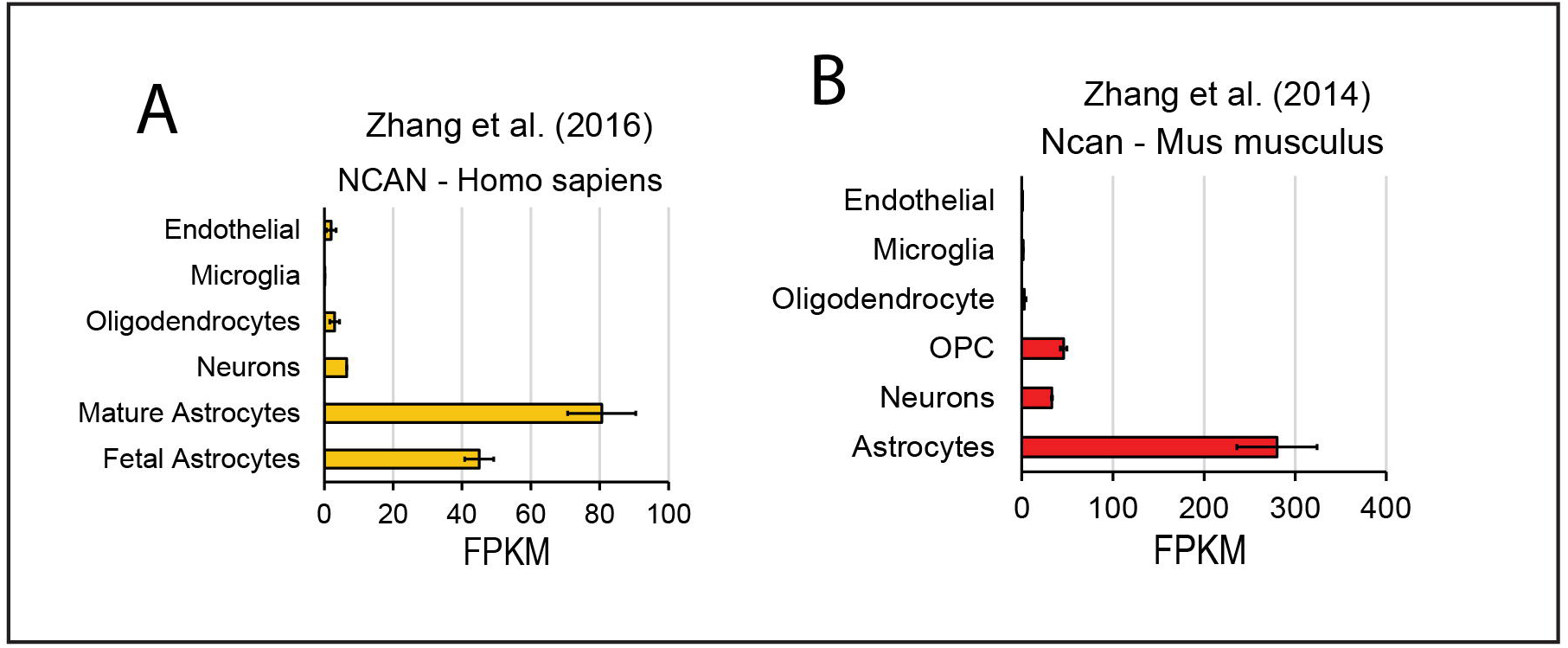
NCAN is highly expressed by human and mouse astrocytes in vivo. Related to Figure 2. (A and B) NCAN expression from RNA-Seq performed on different cell types isolated from mouse and human brain. Human data - Zhang et al. (2016). Mouse data - Zhang et al. (2014).

**Figure S3:**
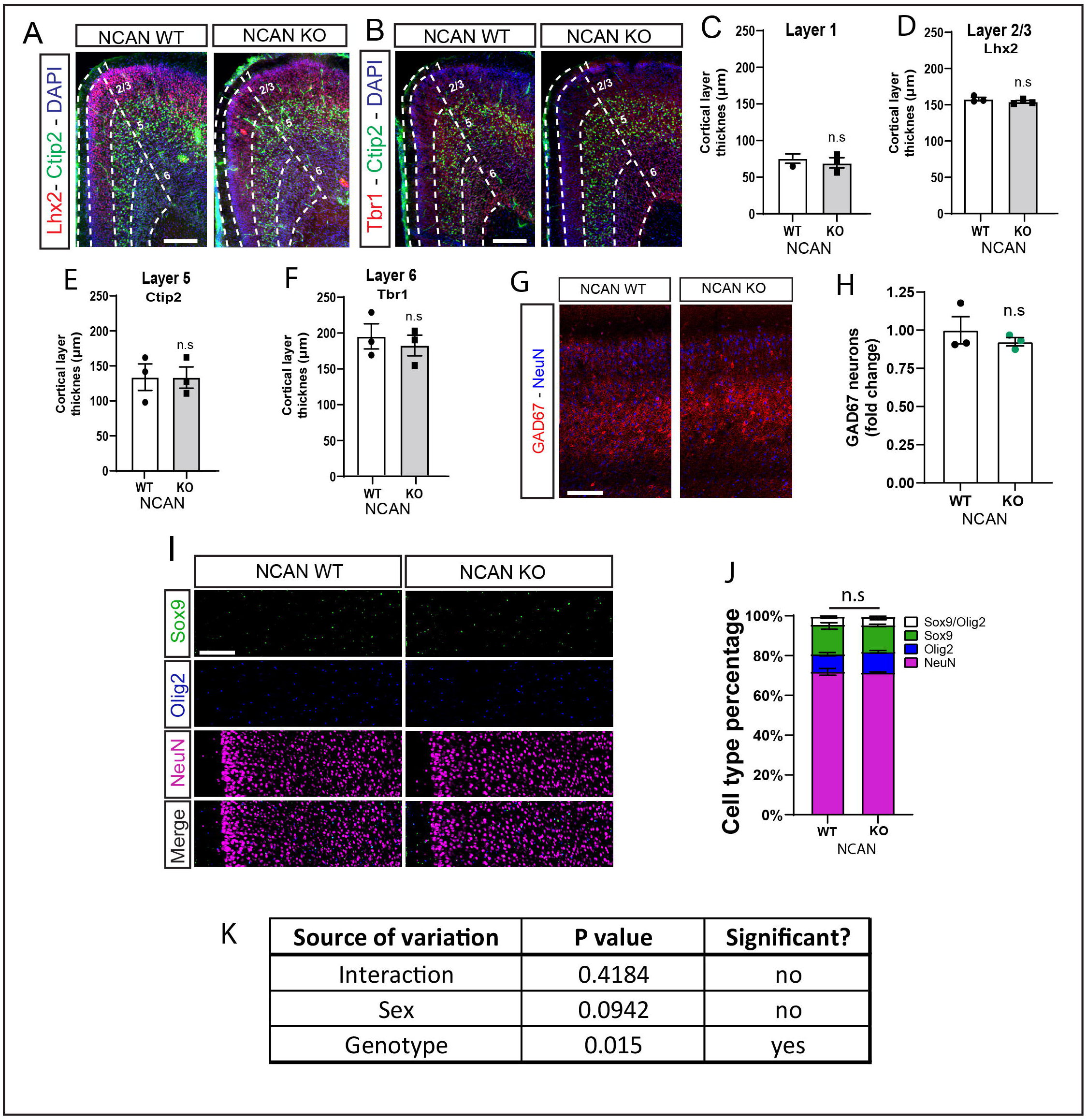
Deletion of NCAN does not alter cortical layer thickness or cell number in the cortex. Related to Figure 3. (A and B) Representative images of the ACC in P1 NCAN WT or KO mice stained with DAPI and neuronal layer markers Lhx2 (layer2-3), Ctip2 (Layer 5) and Tbr1 (Layer 6). Scale bar: 150μm. (C-F) Quantification of cortical thickness in layer 1 (C), layer 2-3 (Lhx2, D), layer 5 (Ctip2, E) and layer 6 (Tbr1, F) of P1 NCAN WT and KO mice. 3 sections/mouse, 3 sex-matched littermate pairs. Data points represent mouse averages. Bars are mean ± s.e.m. Unpaired two-tailed t-test. (G) Representative images of the ACC at P30 in NCAN WT and KO mice stained with GAD67 and NeuN. Scale bar: 100μm. (H) Quantification of GAD67 positive interneurons in NCAN WT and KO mice in the ACC at P30. 3 sections/mouse, 3 sex-matched littermate pairs. Data points represent mouse averages. Bars are mean ± s.e.m. Unpaired two-tailed t-test. (I) Representative images of P30 ACC from NCAN WT and KO mice showing cells Sox9+ (astrocytes), Olig2+ (oligodendrocytes) and NeuN+ (neurons). Scale bar: 75 μm. (J) Quantification of cell number based on nuclear markers. n = 3 sex-matched littermate groups. Bars are mean ± s.e.m. Unpaired two-tailed t-test. (K) Two-way ANOVA analysis of the synapse numbers in WT and littermate NCAN KO mice show only a significant effect of genotype on synapse numbers, no significant effect of sex, and no interaction between sex and genotype. Data from NCAN WT or KO mice in layer 2/3 of the ACC. 5 images/section, 3 sections/mouse, 6 sex-matched littermate pairs.

**Figure S4:**
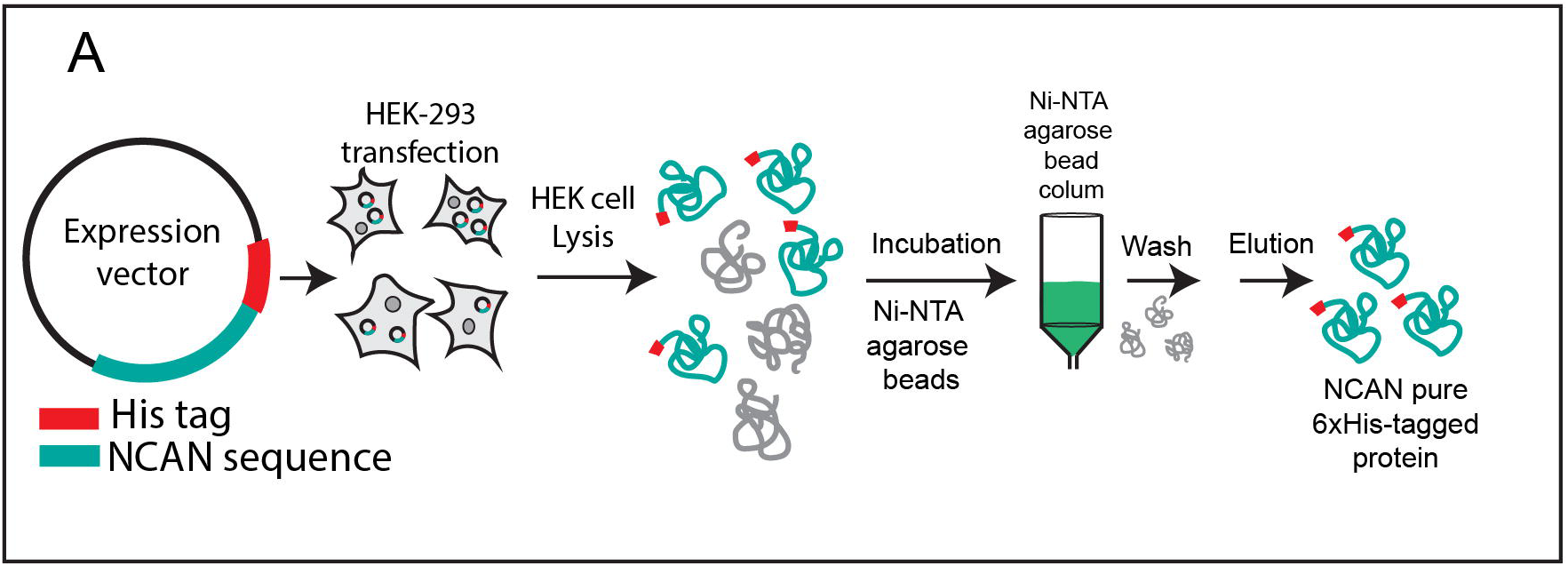
NCAN recombinant protein production and isolation. Related to Figure 4. **(A)** Ni-NTA agarose beads were used to purify NCAN recombinant proteins containing a polyhistidine (6xHis) sequence. Transfected HEK-293 expressed the overexpression constructs. After cell lysis the recombinant proteins bind to the beads and can be eluted by competition with imidazole.

**Figure S5:**
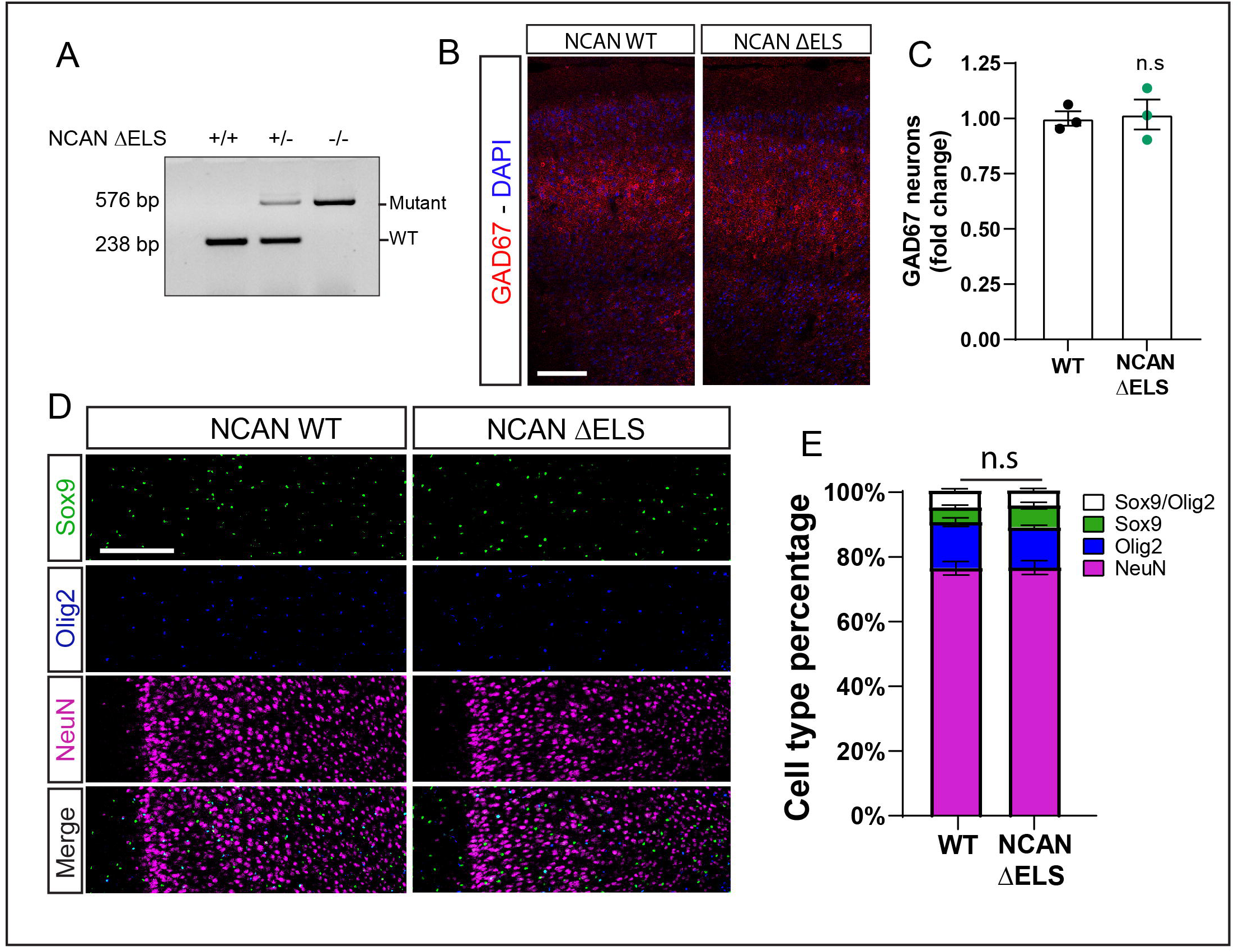
Deletion of NCAN ΔELS domain does not alter cell number in the cortex. Related to Figure 5. (A) Genomic PCR from NCAN WT, Het and ΔELS mutant mice. Mutant band: 576bp, WT band: 238bp. (B) Representative images of the ACC at P30 in NCAN WT and ΔELS mutant mice stained with GAD67 and NeuN. Scale bar: 100μm. (C) Quantification of GAD67 positive interneurons in NCAN WT and ΔELS mutant mice in the ACC at P30. 3 sections/mouse, 3 sex-matched littermate pairs. Data points represent mouse averages. Bars are mean ± s.e.m. Unpaired two-tailed t-test. (D) Representative images of P30 ACC from NCAN WT and ΔELS mutant mice showing cells Sox9+ (astrocytes), Olig2+ (oligodendrocytes) and NeuN+ (neurons). Scale bar: 100 μm. (J) Quantification of cell number based on nuclear markers. n = 3 sex-matched littermate groups. Bars are mean ± s.e.m. Unpaired two-tailed t-test.

**Figure S6:**
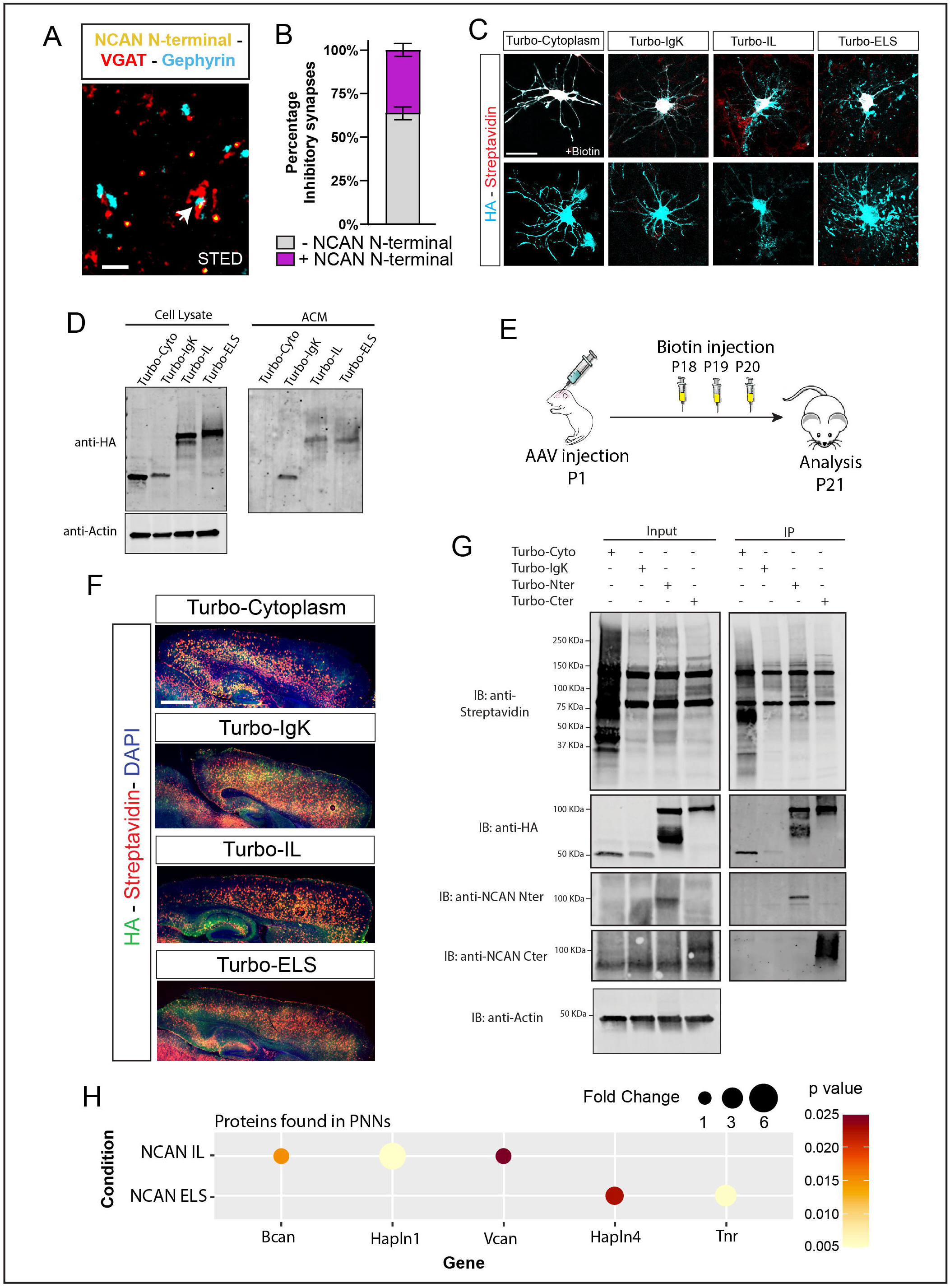
Extracellular Turbo-ID constructs expression and biotinylation *in vitro* and *in vivo*. Related to Figure 6. (A) Three-color STED image of NCAN N-terminal together with VGAT and Gephyrin. Scale bar: 1μm. (B) Percentage of NCAN N-terminal colocalizing with inhibitory synapses. 3 sections/mouse, 3 WT CD1 mice. Bars are mean ± s.e.m. (C) Astrocytes *in vitro* transfected with TurboID constructs (marked with HA), with or without biotin (labeled with streptavidin). Scale bar: 30μm. (D) Western blot analysis of TurboID constructs expression *in vitro* on astrocytes and secretion to the ACM. (E) Diagram of incracortical delivery of TurboID adenovirus injected at P1 in CD1 WT mice. Biotin injections performed at P18, P19 and P20. Brains were harvested and processed at P21 for proteomic analysis. (F) Representative images of *in vivo* expression in the cortex of different TurboID constructs labeled with HA and the biotinylating activity labeled with streptavidin. Scale Bar: 300μm. (G) Western blot analysis of TurboID constructs expression (HA) and biotinylation activity (Streptavidin) *in vitro* in cortical lysates and subsequent immunoprecipitation. (H) Dot plot illustrating the enrichment of different perineuronal net proteins in NCAN IL-TurboID and NCAN ELS-TurboID.

**Figure S7:**
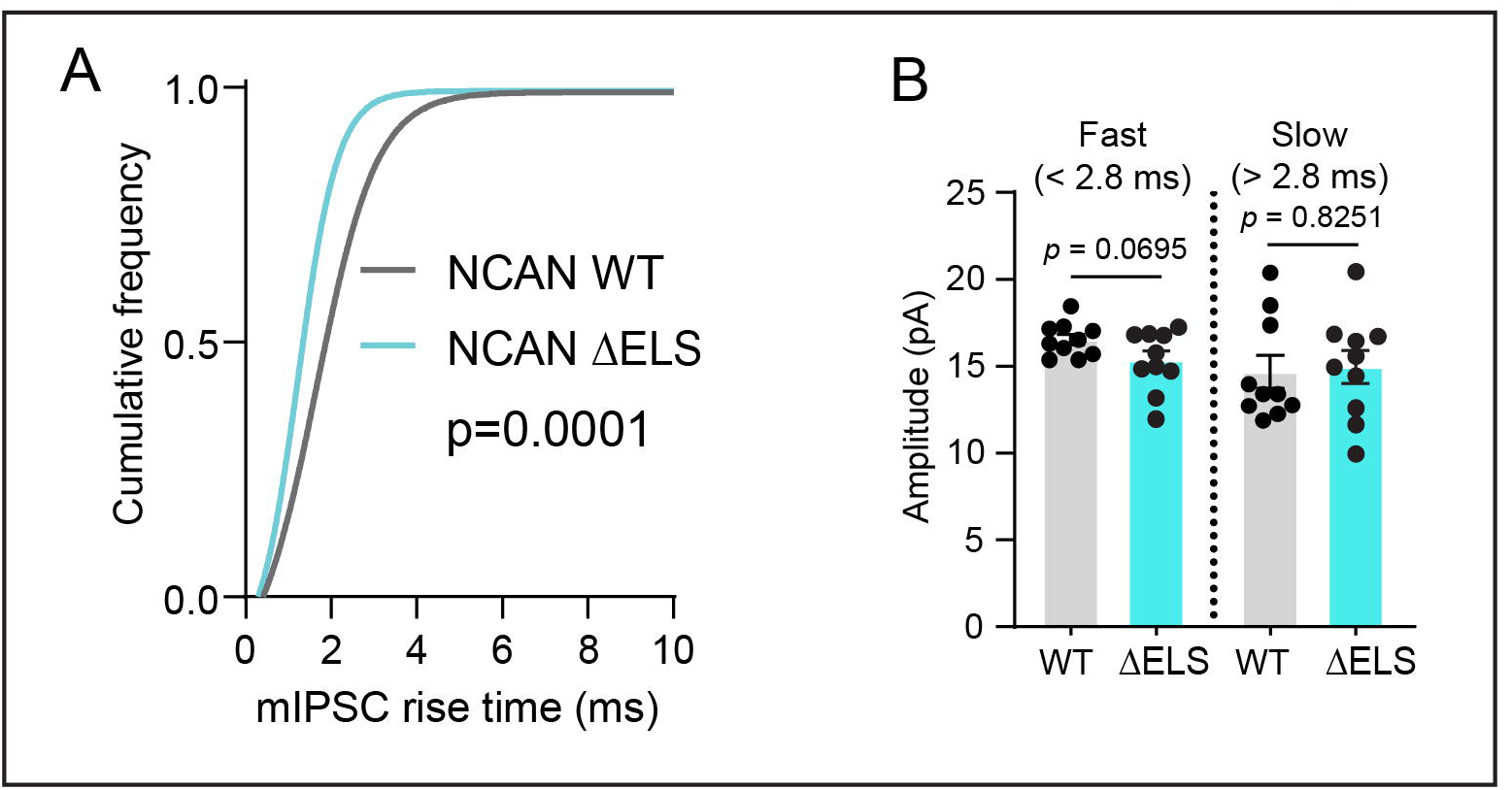
NCAN ΔELS mutant mice electrophysiological recordings differ from WT mice. Related to Figure 7. (A) Quantification of cumulative frequency of mIPSC rising time in P30-35 NCAN WT and ΔELS mutant littermates. n = 10 WT and 10 ΔELS neurons from 3 mice per genotype. Kolmogorov-Smirnov test (D = 0.2647, *p* < 0.0001). (B) Quantification of fast (somatic) and slow (dendritic) amplitude frequency in P30-35 NCAN WT and ΔELS mutant littermates. n = 10 WT and 10 ΔELS neurons from 3 mice per genotype. Average amplitude of fast somatic events in NCAN WT (16.53 ± 0.3077) and ΔELS mutant (15.31 ± 0.5509) mice. Unpaired Two tailed t-test [t (18) = 1.930, *p* = 0.0695]. Average amplitude of slow dendritic events in NCAN WT (14.66 ± 0.9401) and ΔELS mutant (14.96 ± 0.9499) mice. Unpaired Two tailed t-test [t (18) = 0.2242, *p* = 0.8251]. Data are mean ± s.e.m.

## MATERIALS AND METHODS

### KEY RESOURCES TABLE

| REAGENT or RESOURCE | SOURCE | IDENTIFIER |
| --- | --- | --- |
| Actin | Sigma-Aldrich | Cat# A5441, RRID: AB_476744 |
| Bassoon | Enzo/Assay Designs | Cat# ADI-VAM-PS003F, RRID: AB_11181058 |
| Ctip2 | Abcam | Cat# Ab18465, RRID: AB_2064130 |
| GAD-67 | Millipore | MAB5406, RRID:AB_2278725 |
| Gephyrin | Synaptic Systems | Cat# 147 002, RRID:AB_2619838 |
| Gephyrin | Synaptic Systems | Cat# 147 011, RRID:AB_887717 |
| GFAP | Sigma | Cat# G3893, RRID:AB_477010 |
| GFP | Aves labs | GFP-1020, RRID:AB_10000240 |
| HA | Roche | 11867423001, RRID:AB_390918 |
| Homer1 | Synaptic systems | Cat# 160 002, RRID: AB_2120990 |
| Homer1 | Synaptic systems | Cat#160011, RRID: AB_2120992 |
| L1 (ASCS4) | DSHB | Cat# ascs4, RRID:AB_528349 |
| LHX2 | Millipore | Cat# ABE1402, RRID: AB_2722523 |
| NCAN C-terminal | Invitrogen | Cat# PA5-79718, RRID: AB_2746833 |
| NCAN N-terminal | R&D Systems | Cat#AF5800, RRID: AB_2149717 |
| NeuN (A60) | Millipore | Cat# MAB377, RRID:AB_2298772 |
| Olig2 (211F1.1) | Millipore | Cat# MABN50, RRID:AB_10807410 |
| Somatostatin | Millipore | Cat# MAB354, RRID: AB_2255365 |
| Sox9 | Millipore | Cat# AB5535, RRID:AB_2239761 |
| Synaptotagmin-2 | Synaptic Systems | Cat# 105 225, RRID: AB_2744654 |
| Synaptotagmin-2 | Synaptic Systems | Cat# 105 223, RRID: AB_10894084 |
| TBR1 | Abcam | Cat#31940, RRID: AB_2200219 |
| VGAT | Synaptic Systems | Cat# 131 004, RRID:AB_887873 |
| VGluT1 | Millipore | Cat# AB5905, RRID:AB_2301751 |
| WFA | Vector Laboratories | Cat# B-1355 |
| **Bacterial and virus strains** |  |  |
| One Shot STBL3 | Thermo Fisher | Cat# C737303 |
| **Chemicals, peptides, and recombinant proteins** |  |  |
| AraC | Sigma | Cat# C1768 |
| B27 | GIBCO | Cat# 17504044 |
| B27 Plus | GIBCO | Cat# A3582801 |
| BDNF | PeproTech | Cat# 450-02 |
| Carboxypeptidase E | R&D | Cat# 3587-ZN |
| CNTF | PeproTech | Cat# 450-13 |
| Clusterin | R&D | Cat# 2937-HS |
| DMEM | GIBCO | Cat# 11960 |
| DPBS | GIBCO | Cat# 14287 |
| Dystroglycan | R&D | Cat# 6868-DG |
| Fetal Bovine Serum | Thermo Fisher | Cat# 10-437-028 |
| Forskolin | Sigma | Cat# F6886 |
| Glycerol | Acros Organics | Cat# AC158920010 |
| L-Glutamine | GIBCO | Cat# 25030-081 |
| Hydrocortisone | Sigma | Cat# H0888-5G |
| Insulin | Sigma | Cat# I1882 |
| Neurobasal | GIBCO | Cat# 21103049 |
| Neurobasal Plus | GIBCO | Cat# A3582901 |
| Neurobasal minus phenol red | GIBCO | Cat# 12348017 |
| Neurocan Full-length | This paper | N/A |
| Neurocan ELS | This paper | N/A |
| Neurocan IL | This paper | N/A |
| Odyssey Blocking Buffer | LI-COR Biosciences | Cat# 927-40000 |
| Opti-MEM | Thermo Fisher | Cat# 11058021 |
| Paraformaldehyde | Electron microscopy Sciences | Cat# 19210 |
| Pen/Strep | GIBCO | Cat# 15140 |
| PhosSTOP™ | Roche | Cat# 4906845001 |
| Poly-D-Lysine | Sigma | Cat# P6407 |
| Protease Inhibitor Cocktail | Roche | Cat# 4693132001 |
| RIPA Buffer | Sigma | Cat# R0278 |
| Sodium Pyruvate | GIBCO | Cat# 11360-070 |
| DyLight 594 Streptavidin | Vector Laboratories | Cat# SA-5594 |
| TGFβ1 | R&D | Cat# 7666-MB |
| Tissue-Tek® O.C.T. Compound | Sakura® Finetek | Cat# 4583 |
| Triton X-100 | Roche | Cat# 11332481001 |
| **Critical commercial assays** |  |  |
| Endo-Free Maxi Prep Kit | QIAGEN | Cat# 12362 |
| **Experimental models: cell lines** |  |  |
| HEK293T | ATCC | CRL-11268 |
| Rat primary cortical neurons | This paper | N/A |
| Rat primary cortical astrocytes | This paper | N/A |
| Mouse primary cortical astrocytes | This paper | N/A |
| **Experimental models: organisms/strains** |  |  |
| Mouse: Aldh1L1-EGFP | MMRRC | RRID:MMRRC_011015-UCD |
| Mouse: CD1 | Charles River | RRID:IMSR_CRL:022 |
| Mouse: Neurocan KO | This paper | N/A |
| Mouse: Neurocan ΔELS | This paper | N/A |
| Rat: Sprague-Dawley | Charles River | 001 |
| **Oligonucleotides** | Available in Table S4 |  |
| **Recombinant DNA** |  |  |
| pZac2.1-GfaABC1D -IgK-TurboID- NCAN IL-HA | This paper | N/A |
| pZac2.1-GfaABC1D -IgK-TurboID- NCAN ELS-HA | This paper | N/A |
| pZac2.1-GfaABC1D -IgK-TurboID-HA | This paper | N/A |
| pZac2.1-GfaABC1D -TurboID-HA | Takano et al, 2020 | N/A |
| pAPtag5-IgK-NCAN-Full length | This paper | N/A |
| pAPtag5-IgK-NCAN ELS | This paper | N/A |
| pAPtag5-IgK--NCAN IL | This paper | N/A |
| **Software and algorithms** |  |  |
| GraphPad Prism 8 | GraphPAD | https://www.graphpad.com/scientific-software/prism/ |
| ImageJ | NIH | https://imagej.nih.gov/ij/ |
| Puncta Analyzer | Savage et al, 2023 | https://github.com/Eroglu-Lab/Syn_Bot |
| Image Studio | LICOR | https://www.licor.com |
| **Other** |  |  |
| Digital restoration and image segmentation | This paper | https://github.com/ErogluLab/CellCounts |

### RESOURCE AVAILABILITY

#### Lead contact

Further information and requests for resources and reagents should be directed to and will be fulfilled by the lead contacts, Dolores Irala and Cagla Eroglu.

#### Materials availability

The reagents generated in this study are available without restriction.

#### Data and code availability

The data generated during this study are available from the lead contact upon request.

### EXPERIMENTAL MODEL AND SUBJECT DETAILS

#### Animals

All rodents in this study were used in accordance with the Institutional Animal Care and Use Committee (IACUC) and the Duke Division of Laboratory Animal Resources (DLAR) oversight (IACUC Protocol Numbers A147-17-06 and A117-20-05). All mice were housed under typical day/night conditions of 12-hour cycles. Aldh1L1-EGFP (RRID:MMRRC_011015-UCD) mice were obtained through MMRRC. Wild-type CD1 mice used for immunofluorescence, western blot, and TurboID experiments were purchased through Charles River Laboratories (RRID:IMSR_CRL:022). Neurocan knockout mice and Neurocan ΔELS mutant mice were generated in collaboration with the Duke Transgenic Mouse Facility using CRISPR-EZ strategy developed by Modzelewski et al., 2018^90^. Briefly, single guide RNA (sgRNA) was designed and then generated by in-vitro transcription of a short DNA template. After donor superovulation, pronuclear-stage embryos were collected, cultured, and electroporated with the sgRNA and Cas9 Ribonucloprotein (IDT, Alt-R Cas9 v3 Cat#1081059). Viable embryos were cultured and genotyped *ex vivo* to test the sgRNA efficiency. Successful embryos were oviduct transferred to pseudopregnant CD1 females. After birth, funders were screened by PCR with genomic DNA. PCR products were subcloned into pL253 at the NotI-SpeI restriction sites. From each funder, independent subclones were sequenced to identify the targeted mutation. P1, P7, P10, P15, P21 and P30 mice were used for experiments or as specified in the text and figure legends. Littermate pairs of the same sex were used for all the experiments and were randomly assigned to experimental groups based on genotype. Mice of both sexes were included in the analysis of each experiment, and no influence or association of sex on the experimental outcomes was observed for any of the results.

### PRIMARY CULTURES

#### Cortical neuron isolation and culture

Postnatal glia-free rat cortical neuronal cultures were prepared from male P1 rat pups (Sprague Dawley, Charles River Laboratories, SD-001). Cerebral cortices were micro-dissected and enzymatically dissociated with papain (~7.5 units/mL) at 33°C for 45 minutes. After incubation, the cortices were mechanically dissociated in low and high ovomucoid solutions to obtain a single-cell suspension. Next the cells were resuspended in panning buffer (DPBS (GIBCO 14287) supplemented with BSA and insulin) and passed through a 20 μm mesh filter (Elko Filtering 03-20/14) to remove any debris. The filtered cells were incubated on negative panning dishes coated with Bandeiraea Simplicifolia Lectin 1 (x2), followed by goat anti-mouse IgG+IgM (H+L) (Jackson ImmunoResearch 115-005-044), and goat anti-rat IgG+IgM (H+L) (Jackson ImmunoResearch 112-005-044) coated plates. Finally, the cells were incubated for 45 minutes on positive panning dishes coated with mouse anti-L1 (ASCS4, Developmental Studies Hybridoma Bank, Univ. Iowa) to bind only cortical neurons. After incubation, the positive plates were washed with panning buffer and the remaining adherent cells were collected by forceful pipetting with a P1000 pipette. Isolated neurons were pelleted (11 min at 200 g) and resuspended in serum-free neuron growth media (NGM; Neurobasal, B27 supplement, 2 mM L-Glutamine, 100 U/mL Pen/Strep, 1 mM sodium pyruvate, 4.2 μg/mL Forskolin, 50 ng/mL BDNF, and 10 ng/mL CNTF). 70,000 neurons were plated onto 12 mm glass coverslips coated with 10 μg/mL poly-D-lysine (PDL, Sigma P6407) and 2 μg/mL laminin. The neurons were incubated at 37°C in 10% CO_2_. On day *in-vitro* (DIV) 2, half of the media was replaced with NGM Plus (Neurobasal Plus, B27 Plus, 100 U/mL Pen/Strep, 1 mM sodium pyruvate, 4.2 μg/mL Forskolin, 50 ng/mL, BDNF, and 10 ng/mL CNTF) supplemented with AraC (4 μM). On DIV 3, all of the media was replaced with NGM Plus to remove any traces of AraC. At DIV 6 the neurons were fed with NGM Plus. On DIV 8 and DIV 11 the neurons were fed with NGM Plus with ACM or recombinant proteins as indicated on each experiment.

#### Cortical astrocyte isolation and culture

Rat and mouse cortical astrocytes were prepared as described previously^45^. Briefly, P1 rat or mouse cortices from both sexes were micro-dissected, enzymatically dissociated with papain, mechanically dissociated in low and high ovomucoid solutions, and resuspended in astrocyte growth media (AGM: DMEM (GIBCO 11960), 10% FBS, 10 μM, hydrocortisone, 100 U/mL Pen/Strep, 2 mM L-Glutamine, 5 μg/mL Insulin, 1 mM Na Pyruvate, 5 μg/mL N-Acetyl-L-cysteine). Between 20 million cells were seeded on 75 mm2 flasks (non-ventilated cap) coated with poly-D-lysine and incubated at 37°C in 10% CO2. On DIV 3, non-astrocyte cells were removed by forcefully shaking the flasks until only an adherent monolayer of astrocytes remained. AraC was added to the media from DIV 5 to DIV 7 to eliminate contaminating fibroblasts. On DIV 7, astrocytes were trypsinized (0.05% Trypsin-EDTA) and plated into 10cm dishes with AGM. For astrocyte-conditioned media production, on DIV 8 the media was replaced with minimal media (Neurobasal medium, minus phenol red, 100 U/mL Pen/Strep, 2 mM L-Glutamine, 1 mM Na Pyruvate), and the astrocytes were incubated for 4 days. At DIV 12 the astrocyte-conditioned media was collected, filtered, and concentrated using Vivaspin 20, 5kDa MWCO. For astrocyte transfection, at DIV 8 the astrocytes were transfected with the expression plasmids using Lipofectamine LTX with Plus Reagent (Thermo Scientific) per the manufacturer’s protocol.

### CELL LINES

#### HEK293T cells

HEK293 cells were used to produce adeno-associated virus and recombinant proteins. Cells were cultured in DMEM supplemented with 10% fetal bovine serum, 100 U/mL Pen/Strep, 2 mM L-Glutamine, and 1 mM sodium pyruvate. The cells were incubated at 37°C in 5% CO2 and passaged every 2-3 days.

### SHRNA AND CDNA PLASMIDS

#### NCAN expression constructs

To produce His-tag NCAN Full-length, NCAN IL-domain, and NCAN-ELS DNA plasmids gene block fragments that codify for those sequences were produced in the Duke Transgenic Mouse Facility. Each gene block fragment was inserted into a vector backbone pAPtag5 using XbaI and NheI enzyme sites. The secretion signal peptide IgK leader sequence was inserted in the N-terminal of each gene block to promote the equal secretion of the resulting proteins. A 6X His-tag was added to the C-terminal of each fragment for detection. All constructs were confirmed by DNA sequencing

#### TurboID expression constructs

Cytosolic pZac2.1_GFABC1D-Turbo-BirA-HA construct was a gift from Soderling Lab^56^. IgK-TurboID, IgK-TurboID-NCAN IL-HA and, IgK-TurboID-NCAN ELS-HA were cloned into pZac2.1_GFABC1D-Turbo-BirA-HA construct from Takano et al., 2020. To generate these constructs gene block fragments that codify for those sequences were produced in the Duke Transgenic Mouse Facility and cloned into the backbone using EcoRI-HF Restriction Enzyme (NEB) and In-Fusion HD Cloning Kit (Takara) following the manufacturer’s protocol.

#### His-tagged protein purification

For NCAN Full-length, NCAN IL-domain, and NCAN-ELS His-tag protein purification, DNA plasmid of each condition was transfected to 10cm dishes of HEK cells. Two days after transfection the HEK cells were washed with PBS and conditioned with DMEM minimal media (DMEM medium minus phenol red, 100 U/mL Pen/Strep, 2 mM L-Glutamine, 1 mM Na Pyruvate) for four days. Next, the conditioned medium was collected and concentrated using Vivaspin 20, 5kDa MWCO. After concentration, the medium was mixed with Ni-NTA resin (Qiagen) aand incubated overnight at 4°C while gently nutating. Next the Ni-NTA Agarose beads were washed with DPBS then collected in a Poly-Prep Chromatography column (Thermo Fisher). NCAN Full-length, NCAN IL-domain, and NCAN-ELS His-tag peptide fragments were eluted by competition with 250 mM Imidazole in DPBS. The final concentration was measured using BCA protein assay kit (Thermo) per manufacturer’s protocol.

### IMMUNOCYTOCHEMISTRY

#### In vitro

Neuronal or astrocyte cultures on glass coverslips were fixed on DIV 12 with warm 4% PFA for 7 minutes and then washed 3 times with PBS to remove any excess of PFA. The neurons were blocked in a blocking buffer containing 50% normal goat serum (NGS) and 0.2% Triton X-100 for 30 minutes. After removal of the blocking buffer, the cells were incubated overnight at 4°C in primary antibodies diluted in blocking buffer containing 10% NGS. After primary antibody incubation, the neuronal cultures were washed with PBS three times and incubated in Alexa Fluor conjugated secondary antibodies and DAPI (Vector Labs) diluted in a blocking buffer containing 10% NGS for 1 hour at room temperature. Finally, the cells were washed again in PBS three times. The coverslips were mounted onto glass slides (VWR Scientific) with a homemade mounting media (90% Glycerol, 20 mM Tris pH 8.0, 0.5% n-Propyl gallate) and sealed with nail polish. Imaging of single optical sections of synaptic markers or astrocyte markers was acquired using an Olympus FV 3000 microscope with a 60x objective and 2x optical zoom. The imaging was always blind to the experimental condition and individual neurons or astrocytes were selected using DAPI. For synaptic analysis, inhibitory and excitatory synapse numbers were obtained using the ImageJ software SynBot (Savage et al, 2023, https://github.com/Eroglu-Lab/Syn_Bot). The same circular area of interest around each neuron was analyzed. In all cases, a minimum of 3 independent experiments were performed as indicated in the figure legend for each experiment. 20 cells were imaged per condition per experiment.

#### In vivo

Mice were anesthetized with 200 mg/kg tribromoethanol (avertin) and perfused with TBS/Heparin and 4% PFA. Following perfusion, the brains were collected and fixed overnight in 4% PFA solution. Next, the brains were washed with TBS to remove any residual PFA and cryoprotected in 30% sucrose. The brains were frozen and stored at −80 in a solution containing 2 parts 30% sucrose and 1-part O.C.T. (TissueTek). The day before immunostaining the brains were sectioned using a cryostat to obtain floating coronal tissue sections of 40 μm thickness and stored in a 1:1 mixture of TBS/glycerol at −20°C. Sections were washed in 1x TBS containing 0.2% Triton X-100 (TBST) for 10 minutes 3 times and then blocked in 10% NGS diluted in TBST. Following blocking the sections were incubated in primary antibody for 48hs at 4°C with shaking. After primary incubation, sections were washed in TBST for 10 minutes 3 times and incubated in Alexa Fluor conjugated secondary antibodies diluted 1:200 (Life Technologies) with DAPI (1:50,000) for 2 hours at room temperature shaking. After this incubation, the sections were washed with TBST for 10 minutes 3 times and mounted onto glass slides using a homemade mounting media (90% Glycerol, 20 mM Tris pH 8.0, 0.5% n-Propyl gallate) and sealed with nail polish. All the images were acquired on confocal Olympus FV 3000 microscope or confocal Leica Stellaris 8 microscope. For STED microscopy, the samples were prepared as described above with changes in primary and secondary antibodies. Primary antibodies were used more concentrated depending on the antibody, and secondary antibody concentrations were used at 1:100. STED-optimized secondary fluorophores were used: Oregon Green 488, Alexa Fluor 488, Alexa Fluor 594 and abberior STAR RED). STED images were acquired on a Leica SP8 STED microscope using a white light excitation laser, 775 depletion laser (red and far red channels), and 560 depletion laser (green channel). Deconvolution of STED images was performed using Huygens Professional software. The researcher acquiring the images was blinded to the experimental group.

#### Cell counting imaging and analysis

Cell counting was performed as described in Baldwin et al., 2021^68^. Briefly, tile scan images containing the anterior cingulated cortex from P30 Ncan WT, Ncan KO or Ncan ΔELS mutant mice were rapidly acquired on an Olympus FV 3000 using the resonant scanner and 20x objective. For labeled nuclei marked with NeuN, Sox9 or Olig2 nuclear markers were identified using a machine-learning-based method for imaged segmentation (U-Net) ^91^. Full source code for this method is available here: https://github.com/ErogluLab/CellCounts. Co-localization of two nuclear markers was determined in ImageJ. 3 sections per brain from 3 sex-matched littermate groups were analyzed.

For GAD67+ interneuron counting in the ACC, tile scan images from P30 Ncan WT, Ncan KO or Ncan ΔELS mutant mice were acquired on the Olympus FV 3000 using the resonant scanner and 20x objective. The number of cells labeled GAD67 was quantified by hand using the cell counter tool in ImageJ. 3 sections per brain from 3 sex-matched littermate groups were analyzed.

#### Synapse imaging and analysis

Synapse staining, imaging, and acquisition were performed as previously described ^68,92^. Briefly, 40 μm thick coronal sections from *Ncan* WT, *Ncan* KO or *Ncan* ΔELS mutant mice containing the anterior cingulate cortex were used. To label inhibitory synapses VGAT and Gephyrin synaptic markers were used. Images were obtain using a Confocal Olympus FV 3000 inverted microscope. Each image was taken using a 60x magnification objective with 1.64x optical zoom. Each z-stack image contains 15 optical sections spaced 0.34 μm apart. Synapses were identified by the colocalization of pre and postsynaptic puncta. Inhibitory synapse numbers were obtained using the ImageJ software SynBot (https://github.com/Eroglu-Lab/Syn_Bot). 3 sections per brain from 6 sex-matched littermate groups were analyzed.

#### STED analysis

Perineuronal net analysis and synaptic colocalization in *Ncan* WT, *Ncan* KO or *Ncan* ΔELS mutant mice was acquired with 93x objective using 5x zoom at a resolution of 2048 × 2048 pixels, for 5 optical sections spaced 0.15 μm apart. Next, the images were deconvolved using Huygens Professional software and maximum projection images representing 0.6 μm in the z-direction were generated for analysis. NCAN C-terminal puncta localization at synapses was quantified by hand using the cell counter tool in ImageJ. 3 sections per brain from 3 sex-matched WT CD1 littermate groups were analyzed as described in the figure legend.

### PROTEIN EXTRACTION AND WESTERN BLOTTING

Protein was extracted from mouse or rat astrocyte cultures using RIPA buffer with protease inhibitors. Cell lysates were collected, incubated on ice for 10 minutes, and centrifuged at 4°C at high speed for 10 minutes to pellet non-solubilized material. The supernatant was collected and stored at −80°C.

To collect protein from brain cortices, mice P7 or younger were euthanized via rapid decapitation, and mice older than P7 were euthanized using chamber CO_2_ administration. Cortices were micro-dissected and mechanically dissociated using RIPA buffer with protease inhibitor in a dounce homogenizer with Teflon pestle. After getting the homogentate, the samples were collected and incubated at 4°C in a rotor for 15 minutes. Next, the samples were centrifuged at high speed for 10 minutes to pellet non-solubilized material. The supernatant was collected and stored at −80°C.

Pierce BSA Protein Assay Kit (Thermo Fisher) was used to determine protein concentration. The lysates were mixed with 2x Laemmli Sample Buffer (BioRad) containing 5% β-ME and incubated at 95°C for 5 minutes to denature proteins. Samples were loaded into 4%–15% gradient pre-cast gels (Bio-Rad) and run at 100 V for 1 hour. Proteins were transferred at 100 V to PVDF membrane (Millipore) for 2 hours, blocked BSA 3% in TBS 0.001% tween and incubated rocking in primary antibody overnight at 4°C. After overnight incubation, the membranes were washed rocking with TBS 0.001% tween 5 times of 10 minutes, incubated in LI-COR secondary antibodies for one hour, washed rocking with TBS 0.001% tween 5 times of 10 minutes, and imaged on an Odyssey Infrared Imaging system using Image Studio software. <u>Protein expression</u> was quantified using Image Studio Lite software.

### ADENO-ASSOCIATED VIRUS (AAV) PRODUCTION AND ADMINISTRATION

Purified AAV was produced in collaboration with the Duke University Viral Vector Core facility. Briefly, HEK293T cells were co-transfected with 3 plasmids: pAd-DELTA F6 (plasmid No. 112867; Addgene), serotype plasmid AAV PHP.eB, and AAV plasmid (pZac2.1-gfaABC1D-TurboID constructs). After transfection, cells were lysed and the replicated viral vectors were collected, purified and stored at −80°C. The viral titer for each vector was calculated by real-time PCR.

### PROTEOMIC ANALYSIS

#### *In vivo* TurboID protein purification

*In vivo* TurboID experiments were performed as previously described in Takano et al., 2020 with modifications ^56^. CD1 P1 mouse pups were anesthetized by hypothermia and 1µl of each concentrated AAV-TurboID vector was injected bilaterally into the cortex using a Hamilton syringe. Pups were monitored until recovered on a heating pad. At P18, P19 and P20 biotin was subcutaneously injected at 24 mg/kg to increase the biotinylation efficiency. 4 mice of both sexes were used for each TurboID condition. At P21 the cerebral cortices were removed and stored at −80° C. For the protein purification, each cortex was lysed in a buffer containing 50 mM Tris/HCl, pH 7.5; 150 mM NaCl; 1 mM EDTA; protease inhibitor mixture (Roche); and phosphatase inhibitor mixture (PhosSTOP, Roche). Next, an equal volume of buffer containing 50 mM Tris/HCl, pH 7.5; 150 mM NaCl; 1 mM EDTA; 0.4 % SDS; 2 % TritonX-100; 2 % deocycholate; protease inhibitor mixture; and phosphatase inhibitor mixture was added to the samples, following by sonication and centrifugation at 15,000 g for 10 min. The remaining supernatant was ultracentrifuged at 100,000g for 30min at 4° C. Finally, SDS detergent was added to the samples and heated at 45 ° C for 45 min. After cooling on ice, each sample was incubated with High Capacity Streptavidin Agarose beads (ThermoFisher) at 4° C overnight. Following incubation, the beads were serially washed: 1) twice with a solution containing 2% SDS; 2) twice with a buffer 1% TritonX-100, 1% deoxycholate, 25 mM LiCl; 3) twice with 1M NaCl and finally five times with 50 mM ammonium bicarbonate. The biotinylated proteins attached to the agarose beads were eluted in a buffer 125 mM Tris/HCl, pH6.8; 4 % SDS; 0.2 % β-mercaptoethanol; 20 % glycerol; 3 mM biotin at 60°C for 15min.

##### Sample Preparation

The Duke Proteomics and Metabolomics Core Facility (DPMCF) received 16 samples (4 of each NCAN Nter, NCAN Cter, NCAN IGK, and NCAN Cyto) which were kept at −80C until processing. Samples were spiked with undigested bovine casein at a total of either 1 or 2 pmol as an internal quality control standard. Next, samples were supplemented with 10.6 μL of 20% SDS, a final concentration of 1.2% phosphoric acid and 580 μL of S-Trap (Protifi) binding buffer (90% MeOH/100mM TEAB). Proteins were trapped on the S-Trap micro cartridge, digested using 20 ng/μL sequencing grade trypsin (Promega) for 1 hr at 47C, and eluted using 50 mM TEAB, followed by 0.2% FA, and lastly using 50% ACN/0.2% FA. All samples were then lyophilized to dryness. Samples were resolubilized using 120 μL of 1% TFA/2% ACN with 12.5 fmol/μL yeast ADH.

##### LC-MS/MS Analysis

Quantitative LC/MS/MS was performed on 3 μL (25% of total sample) using an MClass UPLC system (Waters Corp) coupled to a Thermo Orbitrap Fusion Lumos high resolution accurate mass tandem mass spectrometer (Thermo) equipped with a FAIMSPro device via a nanoelectrospray ionization source. Briefly, the sample was first trapped on a Symmetry C18 20 mm × 180 μm trapping column (5 μl/min at 99.9/0.1 v/v water/acetonitrile), after which the analytical separation was performed using a 1.8 μm Acquity HSS T3 C18 75 μm × 250 mm column (Waters Corp.) with a 90-min linear gradient of 5 to 30% acetonitrile with 0.1% formic acid at a flow rate of 400 nanoliters/minute (nL/min) with a column temperature of 55C. Data collection on the Fusion Lumos mass spectrometer was performed for three difference compensation voltages (−40v, −60v, −80v). Within each CV, a data-dependent acquisition (DDA) mode of acquisition with a r=120,000 (@ m/z 200) full MS scan from m/z 375 – 1500 with a target AGC value of 4e5 ions was performed. MS/MS scans were acquired in the ion trap in Rapid mode with a target AGC value of 1e4 and max fill time of 35 ms. The total cycle time for each CV was 0.66s, with total cycle times of 2 sec between like full MS scans. A 20s dynamic exclusion was employed to increase depth of coverage. The total analysis cycle time for each injection was approximately 2 hours.

##### Quantitative Data Analysis

Following UPLC-MS/MS analyses, data were imported into Proteome Discoverer 2.5 (Thermo Scientific Inc.). In addition to quantitative signal extraction, the MS/MS data were searched against the SwissProt *M. musculus* database (downloaded in Nov 2019) and a common contaminant/spiked protein database (bovine albumin, bovine casein, yeast ADH, etc.), and an equal number of reversed-sequence “decoys” for false discovery rate determination. Sequest with Infernys enabled (v 2.5, Thermo PD) was utilized to produce fragment ion spectra and to perform the database searches. Database search parameters included variable modification on Met (oxidation). Search tolerances were 2ppm precursor and 0.8Da production with full trypsin enzyme rules. Peptide Validator and Protein FDR Validator nodes in Proteome Discoverer were used to annotate the data at a maximum 1% protein false discovery rate based on q-value calculations. Peptide homology was addressed using razor rules in which a peptide matched to multiple different proteins was exclusively assigned to the protein that has more identified peptides. Protein homology was addressed by grouping proteins that had the same set of peptides to account for their identification. A master protein within a group was assigned based on % coverage.

Prior to imputation, a filter was applied such that a peptide was removed if it was not measured in at least 2 unique samples (50% of a single group). After that filter, any missing data missing values were imputed using the following rules: 1) if only one single signal was missing within the group of three, an average of the other two values was used or 2) if two out of three signals were missing within the group of three, a randomized intensity within the bottom 2% of the detectable signals was used. To summarize the protein level, all peptides belonging to the same protein were summed into a single intensity. These protein levels were then subjected to a normalization in which the top and bottom 10 percent of the signals were excluded and the average of the remaining values was used to normalize across all samples. These normalized protein level intensities were used for the remained of the analysis.

### ELECTROPHYSIOLOGY

For whole-cell patch-clamp recordings, 3-4 mice were used to miniature inhibitory postsynaptic current (mIPSC) for each genotype. During all recordings, brain slices were continuously perfused with standard aCSF at RT (~25°C) and visualized by an upright microscope (BX61WI, Olympus) through a 40x water-immersion objective equipped with infrared-differential interference contrast optics in combination with digital camera (ODA-IR2000WCTRL). Patch-clamp recordings were performed by using an EPC 10 patch-clamp amplifier, controlled by Patchmaster Software (HEKA). Data were acquired at a sampling rate of 50 kHz and low-pass filtered at 6 kHz.

To prepare acute brain slices, after decapitation, the brains were immersed in ice-cold artificial cerebrospinal fluid (aCSF, in mM): 125 NaCl, 2.5 KCl, 3 mM MgCl2, 0.1 mM CaCl2, 10 glucose, 25 NaHCO3, 1.25 NaHPO4, 0.4 L-ascorbic acid, and 2 Na-pyruvate, pH 7.3-7.4 (310 mOsmol). Coronal slices containing the ACC were obtained using a vibrating tissue slicer (Leica VT1200; Leica Biosystems). Slices were immediately transferred to standard aCSF (37°C, continuously bubbled with 95% O2 – 5% CO2) containing the same as the low-calcium aCSF but with 1 mM MgCl2 and 1-2 mM CaCl2. After 30 min incubation, slices were transferred to a recording chamber with the same extracellular buffer at room temperature (RT: ~25°C).

To measure mIPSC, the internal solution contained the following (in mM): 77 K-gluconate, 77 KCl, 10 HEPES, 1 EGTA, 4.5 MgATP, 0.3 NaGTP, and 10 Na-phosphocreatine, pH adjusted to 7.2 – 7.4 with KOH and osmolality set to ~ 300 mosM. mIPSCs were measured in the aCSF bath solution containing 1 µM tetrodotoxin and 10 µM 6-cyano-7-nitroquinoxaline-2,3-dione (CNQX) and 50 µM D-2-amino-5-phosphonopentanoate (D-AP5) at −70 mV in voltage-clamp mode. Series resistance was monitored throughout all recordings and only recordings that remained stable over the recording period (≤30 MΩ resistance and <20% change in resistance) were included. mIPSCs recorded at −70 mV were detected and analyzed using Minhee Analysis software (https://github.com/parkgilbong/Minhee_Analysis_Pack)^1^. To analyze the frequency, events were counted over 5 minutes of recording. To obtain the average events for each cell, at least 100 non-overlapping events were detected and averaged. The peak amplitude of the average mIPSC was measured relative to the baseline current and only events larger than 5 pA were included. Rise time was defined as the time from 10–90% of the peak. Rise time analysis was performed using Stimfit software (https://github.com/neurodroid/stimfit)^2^ and events larger than 10 pA were included. All chemicals were purchased from Sigma-Aldrich or Tocris.

### TRANSMISSION ELECTRON MICROSCOPY

For EM analysis of mouse ACC, P30 NCAN WT controls and KO littermates (3 mice per genotype/age) were first anesthetized with 200 mg/kg tribromoethanol (avertin) and transcardially perfused with PBS solution to clear out blood cells, and then with 2% PFA, 2.5% glutaraldehyde, 2 mM CaCl_2_, and 4 mM MgCl_2_ in 0.1 M cacodylate buffer (pH 7.4). 400 µm thick coronal sections per mouse were cut with a blade and the ACC area was dissected out with a scalpel blade. The slices were immersed in 2% glutaraldehyde, 2 mM CaCl_2_, and 4 mM MgCl_2_ in 0.1 M cacodylate buffer (pH 7.4) and fixed overnight at 4°C. At the Duke Electron Microscopy Service core facility, slices were rinsed in 0.1 M phosphate buffer and postfixed in 1% OsO_4_. Next, the samples were dehydrated in ethanol/acetone and incubated in 50:50 acetone:epoxy overnight at room temperature. Ultrathin serial sections were cut and imaged at the Duke Electron Microscopy Service core facility. Inhibitory synapses (symmetric synapses) were identified and counted when presenting a thin postsynaptic density and oval shape presynaptic vesicles. Ten images per brain from 3 sex-matched littermate groups were analyzed.

### QUANTIFICATION AND STATISTICAL ANALYSIS

GraphPAD Prism 8 or 9 was used for all statistical analyses including Student’s t-test and One-way ANOVA followed by followed by post-hoc Tukey’s test or Dunnett’s test when appropriated. Data was plotted as a super plot to visualize individual data points and averages for each experiment when possible. Each figure legend indicates the n for each experiment and the statistical test used. All data are represented as mean ± standard error of the mean. On each graph, the exact *P*-value is indicated in the figure. Sample sizes were determined based on previous experience for each experiment to yield high power to detect specific effects. No statistical methods were used to predetermine the sample size. All animals were healthy at the time of analysis and both males and females were equally included in the analysis of each experiment.

## ACKNOWLEDGMENTS

This work was supported by a Holland-Trice Brain Research Award and the National Institutes of Health grants U19NS123719 and R01NS102237 to C.E. D.I. was supported by postdoctoral fellowships from the Foerster-Bernstein Family and The Pew Charitable Trust Latin American Fellows Program in the Biomedical Sciences. We thank the Duke Light Microscopy Core Facility, the Duke Transgenic Mouse Facility, the Duke Electron Microscopy Service core facility, The Duke Proteomics, and Metabolomics Core Facility, and the Duke University Viral Vector Core facility. C.E. is an HHMI Investigator. This article is subject to HHMI’s Open Access to Publications policy. HHMI lab heads have previously granted a nonexclusive CC BY 4.0 license to the public and a sublicensable license to HHMI in their research articles. Pursuant to those licenses, the author-accepted manuscript of this article can be made freely available under a CC BY 4.0 license immediately upon publication.

## AUTHOR CONTRIBUTIONS

Conceptualization, D.I. and C.E.; Methodology, D.I., S.W., K.S., F.P.U.S., and C.E.; Investigation, D.I., S.W., K.S., F.P.U.S., L.N., and D.S.B; Formal Analysis, D.I., S.W., K.S., F.P.U.S., L.N.; Writing – Original Draft, D.I. and C.E.; Writing – Review & Editing, D.I., S.W., K.S., F.P.U.S., L.N., D.S.B and C.E.; Funding Acquisition, D.I., and C.E.

